# Analysis of endogenous NOTCH1 from *POFUT1 S162L* patient fibroblasts reveals the importance of the *O*-fucose modification on EGF12 in human development

**DOI:** 10.1101/2024.04.09.588484

**Authors:** Kenjiroo Matsumoto, Kelvin B. Luther, Robert S. Haltiwanger

## Abstract

NOTCH1 (N1) is a transmembrane receptor interacting with membrane-tethered ligands on opposing cells that mediate the direct cell-cell interaction necessary for many cell fate decisions. Protein *O*-fucosyltransferase 1 (POFUT1) adds *O*-fucose to Epidermal Growth Factor (EGF)-like repeats in the NOTCH1 extracellular domain, which is required for trafficking and signaling activation. We previously showed that *POFUT1 S162L* caused a 90% loss of POFUT1 activity and global developmental defects in a patient; however, the mechanism by which POFUT1 contributes to these symptoms is still unclear. Compared to controls, *POFUT1 S162L* patient fibroblast cells had an equivalent amount of N1 on the cell surface but showed a 60% reduction of DLL1 ligand binding and a 70% reduction in JAG1 ligand binding. To determine if the reduction of *O*-fucose on N1 in *POFUT1 S162L* patient fibroblasts was the cause of these effects, we immunopurified endogenous N1 from control and patient fibroblasts and analyzed *O*-fucosylation using mass spectral glycoproteomics methods. N1 EGF8 to EGF12 comprise the ligand binding domain, and *O*-fucose on EGF8 and EGF12 physically interact with ligands to enhance affinity. Glycoproteomics of N1 from *POFUT1 S162L* patient fibroblasts showed WT fucosylation levels at all sites analyzed except for a large decrease at EGF9 and the complete absence of *O*-fucose at EGF12. Since the loss of *O*-fucose on EGF12 is known to have significant effects on N1 activity, this may explain the symptoms observed in the *POFUT1 S162L* patient.

## Introduction

The glycosylation of transmembrane receptors plays a crucial role in various steps of cell signaling activation (Moremen, K.W., Tiemeyer, M. and Nairn, A.V. 2012). These glycan modifications can impact protein trafficking (Hammond, C., Braakman, I. and Helenius, A. 1994, Okajima, T., Xu, A., et al. 2005) and enhance or block receptor-ligand interactions (Luca, V.C., Jude, K.M., et al. 2015, Luca, V.C., Kim, B.C., et al. 2017, Wang, J., Miao, Y., et al. 2022). Specifically, *O*-fucose modifications on Notch receptors are crucial for Notch trafficking and signaling activation (Matsumoto, K. and Haltiwanger, R. 2018).

NOTCH1 (N1) is a transmembrane receptor conserved in metazoans that signals through direct cell-to-cell interaction. Differing from most signal receptors, N1 is activated upon ligand binding through regulated intramembrane proteolysis (RIP) resulting in release of the Notch Intracellular Domain (NICD), which translocates to the nucleus where it activates transcription (Kopan, R. and Ilagan, M.X. 2009). N1 is critical in development and its dysregulation or disruption results in numerous developmental disorders (Artavanis-Tsakonas, S. and Muskavitch, M.A. 2010, Bray, S.J. 2006, Kopan, R. and Ilagan, M.X. 2009, Matsumoto, K., Luther, K.B. and Haltiwanger, R.S. 2020, Tien, A.C., Rajan, A. and Bellen, H.J. 2009). In mammalian cells, the activation of Notch signaling requires physical interaction between one of the four NOTCH receptors (N1, NOTCH2 (N2), NOTCH3 (N3), or NOTCH4 (N4)) and one of four signaling ligands (Delta-like1 (DLL1), Delta-like 4 (DLL4), Jagged1 (JAG1), or Jagged2 (JAG2) (Artavanis-Tsakonas, S. and Muskavitch, M.A. 2010).

The extracellular domain of N1 is composed primarily of 36 tandem EGF-like repeats, many of which contain the consensus sequence C^2^-X-X-X-X-S/T-C^3^ for *O*-fucosylation by Protein *O*-fucosyltransferase 1 (POFUT1) (Wang, Y., Shao, L., et al. 2001) (Fig. 1). Previous studies have shown that 21 out of 36 EGF-like repeats in mouse N1 are modified by *O*-fucose (Kakuda, S. and Haltiwanger, R.S. 2017, Matsumoto, K., Kumar, V., et al. 2022). Elimination of *Pofut1* in mice results in embryonic lethality with Notch-like phenotypes (Shi, S. and Stanley, P. 2003). Modification of N1 by POFUT1 has been shown to play a role in the trafficking of N1 to the cell surface (Okajima, T., Xu, A., et al. 2005, Takeuchi, H., Yu, H., et al. 2017) and to affect binding and signaling activation through ligands (Kakuda, S. and Haltiwanger, R.S. 2017, Okajima, T. and Irvine, K.D. 2002, Taylor, P., Takeuchi, H., et al. 2014).

**Fig. 1.**
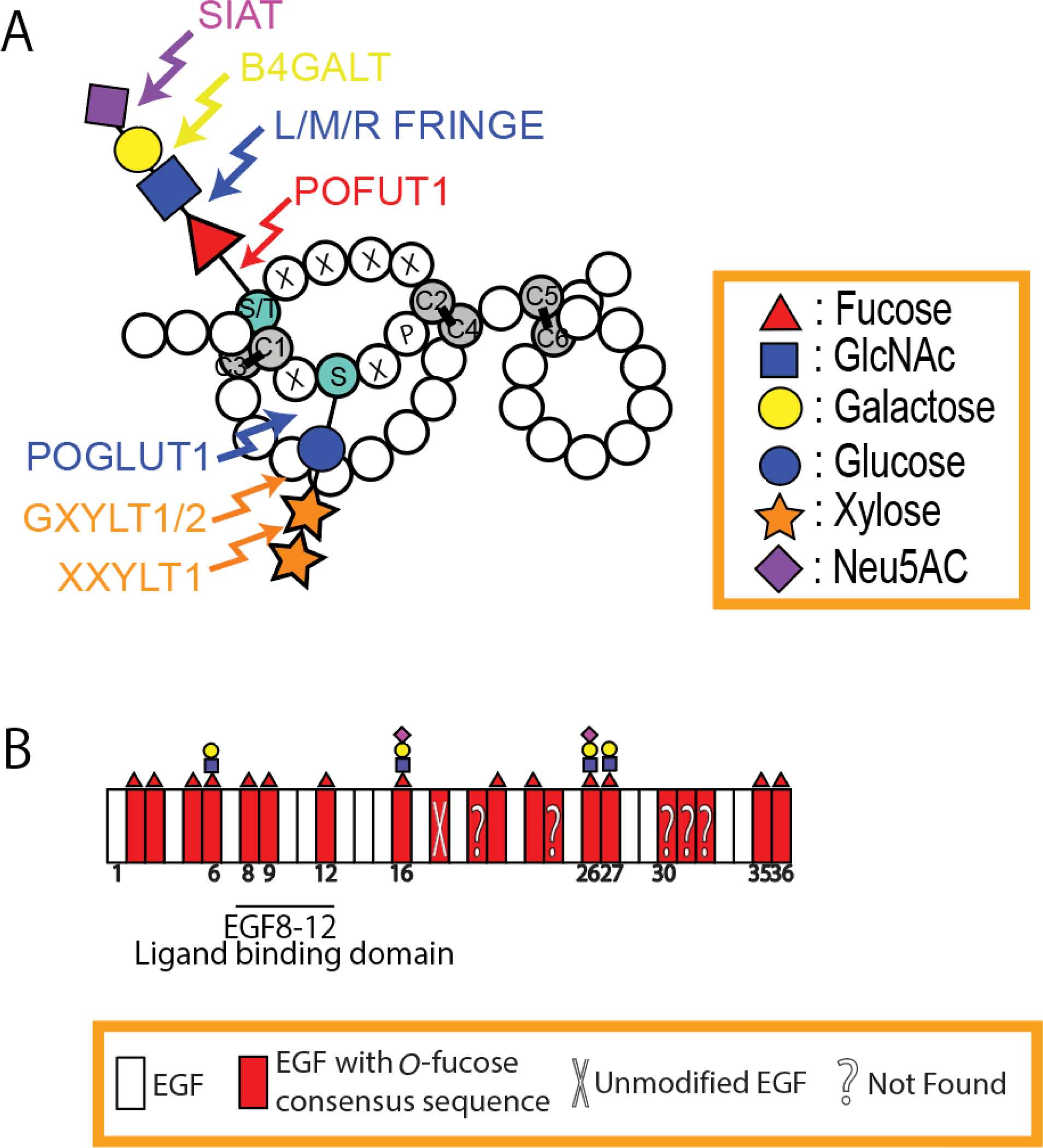
*O*-fucose modifications of NOTCH1. **(A**) Diagram showing an EGF repeat with *O*-fucose and *O*-glucose glycans. Circles are amino acids, gray circles are the conserved cysteines forming disulfide bonds, and consensus sequences for *O*-fucose and *O*-glucose addition are included with single letter amino acid codes. Enzymes responsible for the modifications are indicated. Monosaccharide symbols are based on the Symbol Nomenclature for Glycans (Neelamegham, S., Aoki-Kinoshita, K., et al. 2019). (**B**) Summary of mass spectral analysis of endogenous mouse N1 from activated T cells. The most elongated form of the *O*-fucose glycan detected at each EGF is shown. A question mark indicates that a peptide with the *O*-fucose consensus site in the EGF was not found. Modified from (Matsumoto, K., Kumar, V., et al. 2022) with permission.

N1 EGF8 to EGF12 has been shown to be the ligand binding domain (Luca, V.C., Jude, K.M., et al. 2015, Luca, V.C., Kim, B.C., et al. 2017) and EGF8, EGF9, and EGF12 are modified by *O*-fucose *in vivo* in flies and mice (Harvey, B.M., Rana, N.A., et al. 2016, Matsumoto, K., Kumar, V., et al. 2022). Mutation of the *O*-fucosylation modification site on EGF12 (*EGF12f*) in both *Drosophila* N and mouse N1 resulted in embryonic lethality (Ge, C. and Stanley, P. 2008, Pandey, A., Harvey, B.M., et al. 2019, Varshney, S., Wei, H.X., et al. 2019). Co-crystal structures of a NOTCH fragment and a fragment of its ligands, DLL4 (Luca, V.C., Jude, K.M., et al. 2015) or JAG1 (Luca, V.C., Kim, B.C., et al. 2017) have been reported, and physical interaction of EGF12 *O*-fucose with the ligand has been demonstrated. These results indicate the importance of *O*-fucose on EGF12.

Recently, Takeuchi *et al.,* reported that a patient with a homozygous *POFUT1 S162L* mutation showed reduced N1 signaling with global developmental delay, failure to thrive, severe coarctation of the aorta, ventricular septal defect, constipation, short stature, microcephaly, reduced fat pads, low set and posteriorly rotated ears, wide-spaced nipples, mild hypotonia, and some coagulation factor abnormalities, although no overt bleeding disorder was observed (Takeuchi, H., Wong, D., et al. 2018). It was recently reported that loss of N1 in dermal fibroblasts leads to enhanced healing from diabetic wounds demonstrating the importance of N1 signaling in fibroblasts (Shao, H., Li, Y., et al. 2020). Additionally, induction of cardiomyocyte specialization into ventricular conduction system-like cells *in vitro* is dependent on N1 signaling in cardiac fibroblasts (Ribeiro da Silva, A., Neri, E.A., et al. 2020). Loss of N1 signaling in *POFUT1 S162L* patient cells could be due to a N1 trafficking defect or from loss of Notch-ligand interactions. Here we show that *POFUT1 S162L* patient fibroblasts did not have N1 trafficking defects but had decreased levels of ligand binding. We immunopurified endogenous human N1 from control and patient fibroblasts and analyzed *O*-fucosylation using mass spectral glycoproteomics. While most *O*-fucose sites were modified at or near WT levels, *O*-fucose was absent from EGF12, a critical site for N1 signaling activation.

## Results

### Fibroblasts from a *POFUT1 S162L* patient exhibit no decrease in cell surface expression of N1

*In vitro* glycosyltransferase assays demonstrate the S162L mutation results in a 90% reduction in POFUT1 function (Takeuchi, H., Wong, D., et al. 2018). *POFUT1*-null HEK293T cells display N1 trafficking defects, resulting in a 50% decrease in cell surface N1 levels in *POFUT1*-null HEK293T cells (Takeuchi, H., Yu, H., et al. 2017). Therefore, we investigated whether N1 trafficking in *POFUT1 S162L* patient cells was perturbed.

We first assessed the POFUT1 S162L protein level in the patient fibroblasts and detected a 50% reduction in POFUT1 S162L protein compared to WT (Fig. 2A). Next, we examined the cell surface expression of N1 by Western blotting of *POFUT1 S162L* patient fibroblast cell lysates. N1 undergoes cleavage by Furin protease (S1 cleavage) in the trans-Golgi network, leading to the generation of a Ca^2+^-dependent heterodimer held together by non-covalent bonds which then traffics to the cell surface. Consequently, two bands are detected for N1 in immuno-blots of cell lysate with an anti-N1 intracellular domain antibody: an ∼110 kD band representing the cleaved transmembrane/intracellular domain (Tm/ICD) from the cell surface and an ∼300 kD band representing the immature uncleaved ER form (N1-full). We lysed normal and *POFUT1 S162L* fibroblasts for Western blotting, stained them with N1-ICD and TUB antibodies, and measured the intensity of each band. The ratio of cleaved to uncleaved N1 intensity is similar in control and patient fibroblasts, indicating no accumulation of unprocessed N1 in *POFUT1 S162L* fibroblasts (Fig. 2B). Interestingly, in the *POFUT1 S162L* cells, there is a statistically significant 10-20% increase in total N1 detected (Fig. 2B). We conclude from these data that the *POFUT1 S162L* mutation does not have a major effect on N1 maturation and trafficking to the cell surface.

**Fig. 2.**
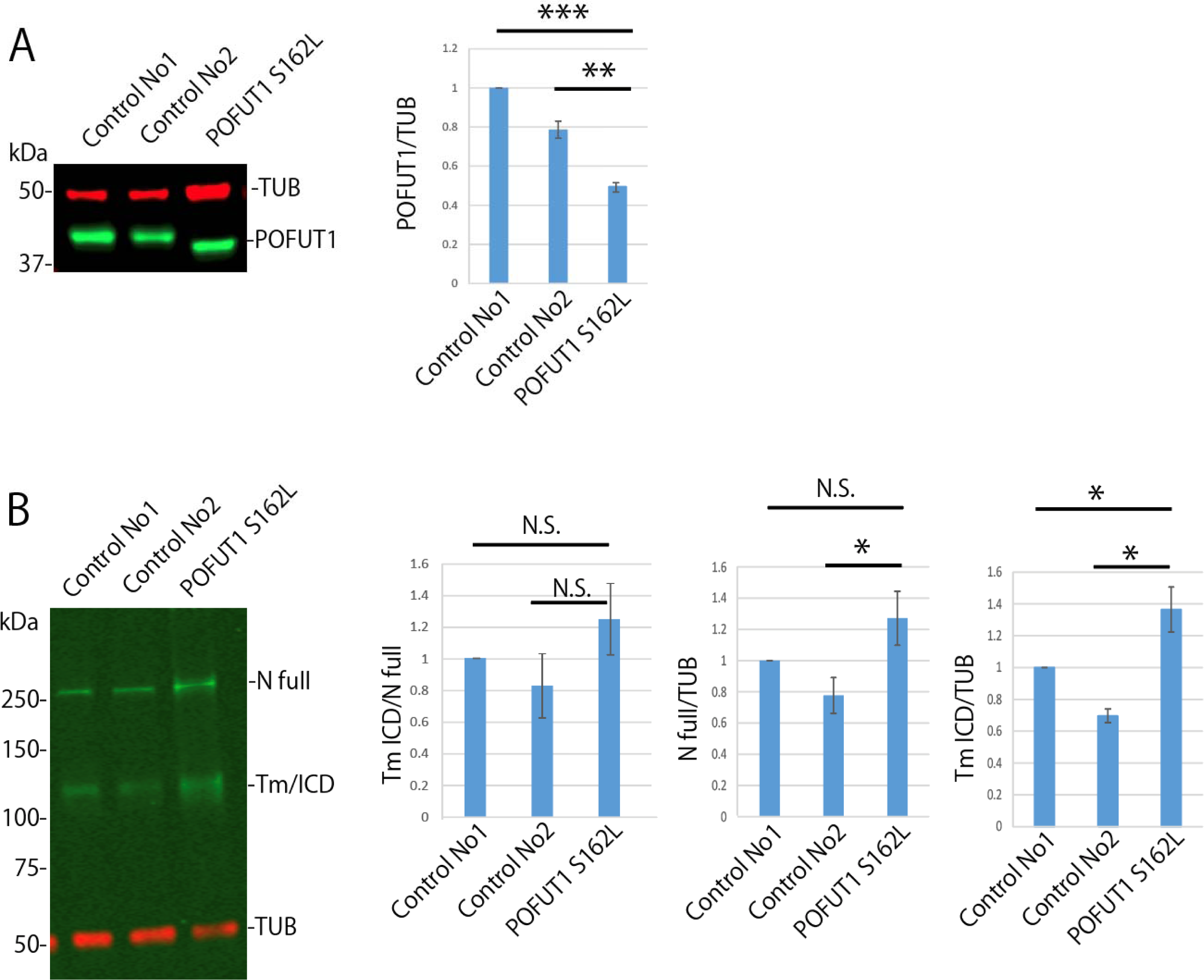
POFUT1 levels are decreased but cell surface N1 is not in *POFUT1 S162L* patient fibroblasts. (**A**) Expression of POFUT1 was evaluated in lysates of control and *POFUT1 S162L* patient fibroblasts by Western blotting stained with anti-POFUT1 antibody and anti-TUBULIN antibody as a control. The intensity of POFUT1 bands was normalized to TUB and shown as bar graphs on the right from biological triplicates. (**B**). Cell surface N1 was checked by Western blotting of control and patient fibroblasts. N1 was detected using a N1 intracellular domain antibody; TUB was used for control. N1 full (above 250 kD) shows N1 accumulated in the ER, while Tm/ICD (above 100 kD) shows cell surface N1. The intensity of N1 full to Tm/ICD bands (left graph), and N1 full or Tm/ICD bands normalized to TUB (two graphs on right) from three biological replicates. *p<0.05, **p<0.01, ***p<0.001. Error bars show +/-SD.

### NOTCH in *POFUT1 S162L* patient cells has a lower affinity for ligands

To investigate the functional status of cell surface NOTCH in *POFUT1 S162L* patient fibroblasts, we conducted cell-based ligand binding assays. In this assay, WT and *POFUT1 S162L* patient fibroblasts were incubated with soluble DLL1-Fc or JAG1-Fc ligands. The presence of Fc was detected using an anti-Fc PE conjugate, and the amount of PE on the cell surface was measured via flow cytometry (Fig. 3). Varying concentrations of DLL1-Fc (0 to 150 nM) (Fig. 3A-C) and JAG1-Fc (0 to 37.5 nM) (Fig. 3E-G) were utilized in each experimental condition. The results are summarized graphically (Fig. 3D and H), revealing a statistically significant 60% reduction in DLL1-Fc binding (Fig. 3D) and a 70% reduction in JAG1-Fc binding (Fig. 3H) in *POFUT1 S162L* patient fibroblasts. Notably, EDTA disrupts the interaction between NOTCH and its ligands (Fig. 3A’**-**C’ and 3E’**-**G’), indicating that these reductions in binding are not a result of nonspecific background staining by the ligand-Fc constructs. These findings suggest that *POFUT1 S162L* patient fibroblasts may have reduced *O*-fucose modification on the ligand binding domain of NOTCH.

**Fig. 3.**
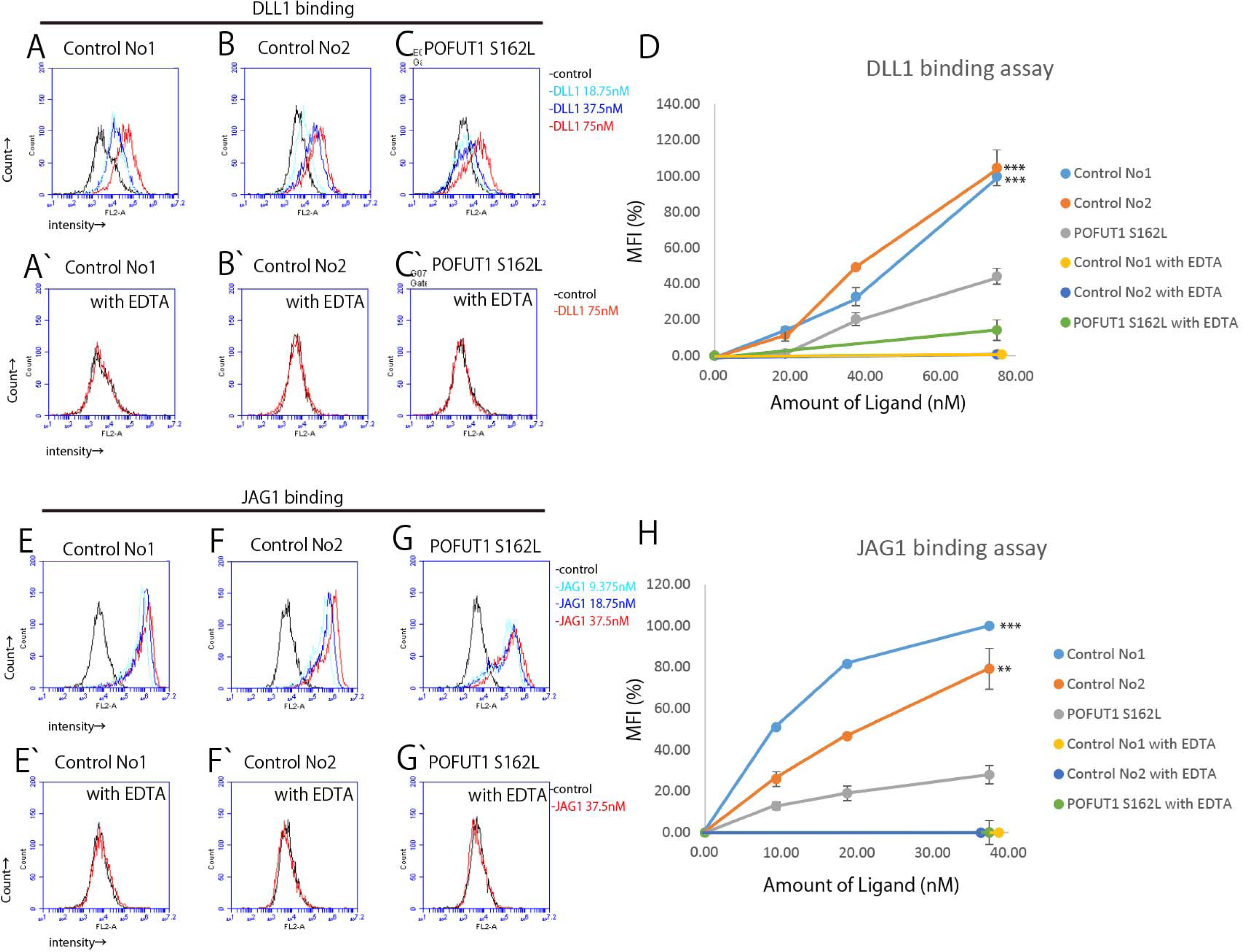
Notch binding to DLL1 or JAG1 is reduced in *POFUT1 S162L* patient fibroblasts Soluble forms of Notch ligands (DLL1-Fc and JAG1-Fc) were used for this ligand binding assay. Control fibroblasts and *POFUT1 S162L* patient fibroblasts were incubated with different concentrations of ligands, stained with anti-Fc PE conjugate, and the cell surface ligands were detected by flow cytometry as described in Materials and Methods. (**A-D**) and (**E-H**) show binding for the indicated amounts of DLL1-Fc and JAG1-Fc, respectively. A’-C’ and E’-G’, indicate the ligand binding assay performed with EDTA, which eliminates Notch-ligand binding. The summary of the results of A-C’ and E-G’ are shown as D and H, respectively. **p<0.01, ***p<0.001. Error bars show +/-SD.

### *O*-fucose modification on EGF12 is lost on N1 in *POFUT1 S162L* patient cells

In our previous work, we developed methods to investigate *O*-fucose and Fringe modifications of mouse N1 immunopurified from activated T cells (Matsumoto, K., Kumar, V., et al. 2022). To analyze endogenous human N1, we identified an antibody capable of efficiently immunopurifying human N1 from HEK293T cells. Fig. S1A shows that a sheep anti-human N1 polyclonal antibody against the extracellular domain (ECD) of human N1 efficiently immunopurified (IPed) N1. The purified human N1 ECD was divided into three portions and digested with trypsin, chymotrypsin, or V8 proteases. The resulting peptides were analyzed by mass spectrometry and the relative level of *O*-fucose modification on each peptide was quantified via extracted ion chromatograms (EICs). Human N1 has 21 predicted *O*-fucose sites, and peptides containing 16 of these sites were identified (Fig. 4A, Fig. S3, Table S1). Peptides containing the *O*-fucose consensus sequence from EGF18, EGF21, EGF30, EGF31, and EGF32 were not detected (Fig. 4B). Searches utilizing Byonic software showed the *O*-fucose monosaccharide modification on all detected peptides with an *O*-fucose consensus site (Fig. 4A, Fig. S3, Table S1). EGF12 and EGF16 had elongated trisaccharide or tetrasaccharide *O*-fucose glycans on a small proportion of *O*-fucosylated peptides (Fig. 4B, Fig. S3). *O*-fucosylation at all sites analyzed was at high stoichiometry except for EGF24 (Fig. 4B).

**Figure 4.**
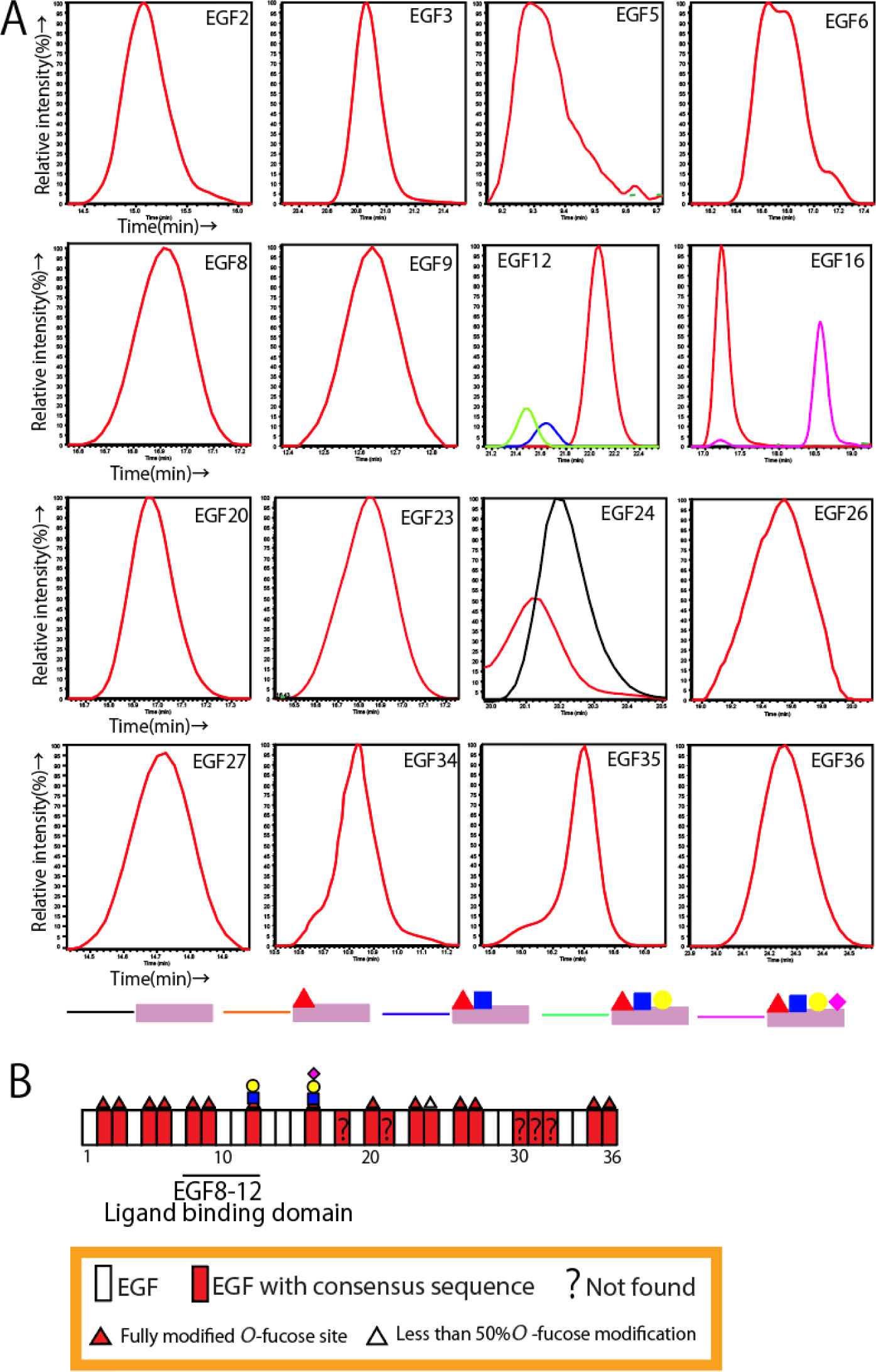
Endogenous N1 from HEK293T cells is modified by *O*-fucose at high stoichiometry. **(A)** EICs showing relative amounts of *O*-fucose peptide glycoforms from the indicated EGFs. Black (unmodified), red (monosaccharide), blue (disaccharide), green (trisaccharide) and purple (tetrasaccharide) lines indicate the curves for the various glycoforms. The EGF12 peptide is also modified with an *O*-glucose trisaccharide (see Fig. S3I). (**B)**. Summary of *O*-fucose modifications identified by mass spectral analysis of endogenous hN1 isolated from HEK293T cells. The most elongated form of the *O*-fucose glycan detected at each EGF is shown. A question mark indicates that a peptide with the *O*-fucose consensus site in the EGF was not found. MS2 spectra for these peptides are in Fig. S3. List of all peptides detected are in Table S1.

Since N1 expression is limited in fibroblasts, obtaining sufficient material for full site mapping is challenging. We therefore targeted our mass spectral analyses mainly to the ligand binding region. Our focus was on EGF12 since the *O*-fucose modification on N1 EGF12 is located within the binding interface of Notch-Delta and Notch-Jagged (Luca, V.C., Jude, K.M., et al. 2015, Luca, V.C., Kim, B.C., et al. 2017) and is known to be one of the essential sites for N1 function (Kakuda, S. and Haltiwanger, R.S. 2017). We purified endogenous N1 from control and *POFUT1 S162L* patient fibroblasts and eluted the protein using 8M urea (Figs. S1B and C). The eluted N1 was subjected to chymotryptic digestion followed by mass spectrometric analysis to examine the level of *O*-fucose modification via EICs (Fig. S2). Interestingly, the *O*-fucose on EGF12 was completely elongated to a trisaccharide in the control sample, suggesting that higher levels of fringe enzymatic activity is present in fibroblasts than in HEK293T cells. The levels of *O*-fucose on the N1 peptides from EGFs 2, 8, 9, 12, and 35 in the EICs in Fig. S2 are summarized in Fig. 5. EGFs at a distance from the ligand binding region such as EGFs 2, and 35 showed stoichiometric levels of fucosylation in both control and *POFUT1 S162L* patient cells (Fig. 5, Fig. S2). All of the EGFs in the ligand binding domain (EGF8-12) were stoichiometrically modified in control fibroblasts. In patient cells, EGF8 had stoichiometric modification, while EGF 9 showed a dramatically decreased level of *O*-fucosylation, and *O*-fucose was completely absent on EGF12 (Fig. 5, Fig. S2). MS2 spectra for each of the EGFs analyzed in Fig. 5 are shown in Fig. S4. These data demonstrate that *POFUT1 S162L* patient cells do not fucosylate EGF12 of N1.

**Fig. 5.**
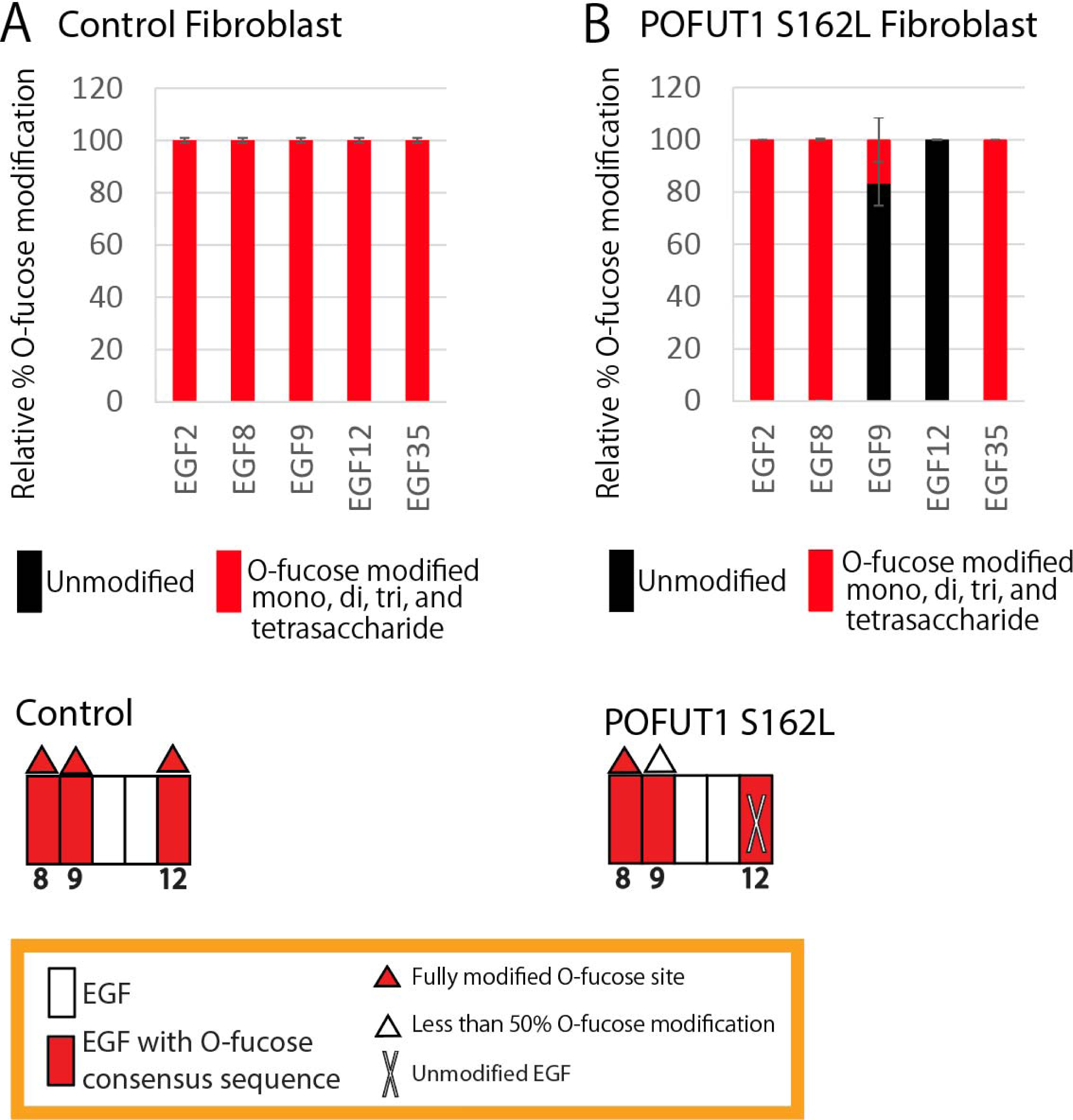
N1 EGF12 is not modified with *O*-fucose in *POFUT1 S162L* patient fibroblasts Top: Bar graphs show the relative amount of *O*-fucose glycoforms on peptides from each EGF repeat detected in N1 from the control fibroblasts **(A)** or *POFUT1 S162L* patient fibroblasts **(B)**. Black and red indicate the unmodified and *O*-fucose-modified peptides, respectively. The average of three biological replicates is shown. Error bars show +/-SD. Representative EICs are shown in Fig. S2. Mass spectral data can be found in Fig. S4 and Table S2. **Bottom:** Summary of *O*-fucose modifications of the ligand binding domain of endogenous N1 from the control fibroblast **(A)** or *POFUT1 S162L* patient fibroblast **(B)**.

## Discussion

The S162L mutation reduces POFUT1 activity more than 90% compared to WT (Takeuchi, H., Wong, D., et al. 2018). To determine the effect of this reduction in POFUT1 activity in patient cells, we examined N1 trafficking, ligand binding, and N1 glycan modification in *POFUT1 S162L* fibroblasts. We observed that maturation and trafficking of N1 was not affected in *POFUT1 S162L* fibroblasts but that both DLL1 and JAG1 binding were significantly reduced. We developed an immunopurification method for endogenous human N1 using HEK293T cells and demonstrated high levels of *O*-fucosylation on many EGFs. We used this same protocol to IP endogenous N1 from control and *POFUT1 S162L* fibroblasts and analyzed the levels of *O*-fucose modification using mass spectral glycoproteomics methods. We observed that EGFs 2, 8 and 35 of endogenous N1 were modified exclusively by monosaccharide *O*-fucose in WT fibroblasts while EGF12 was elongated to a trisaccharide. In *POFUT1 S162L* fibroblasts, *O*-fucosylation was completely absent from the critical EGF12 in the ligand binding region. *O*-fucosylation of EGF9 was also significantly reduced. EGF12 of N1 from *POFUT1 S162L* fibroblasts was however, modified by *O*-glucose trisaccharide (Fig. S4F), a modification that can only be added to properly folded EGFs (Li, Z., Fischer, M., et al. 2017, Takeuchi, H., Kantharia, J., et al. 2012). This demonstrates that the alteration in *O*-fucose modification at this site in patient fibroblasts is not due to folding defects in EGF12 but rather the loss of POFUT1 activity. These findings provide insights into the molecular basis of Notch signaling impairment in *POFUT1 S162L* fibroblasts and highlight the importance of proper *O*-fucose modification for N1 function.

Interestingly, EGF12 contains an aspartic acid residue (D464) in close proximity substrate for POFUT1 than other EGFs with the *O*-fucose consensus sequence. N2 has a D in the same position in EGF12, suggesting it will be poor substrate as well. N3 EGF11 (equivalent of EGF12 in other Notch receptors) and N4 EGF12 do not have a D in the same position, suggesting that these EGFs may be more efficiently modified with *O*-fucose by POFUT1 and thus less sensitive to changes in POFUT1 activity. As with N1, loss of *O*-fucose on N2 EGF12 reduces DLL1 ligand binding (Kakuda, S., LoPilato, R.K., Ito, A., and Haltiwanger, R.S. 2020). Previous studies demonstrated that N1 and N3 are expressed in fibroblasts (Wei, K., Korsunsky, I., et al. 2020). Given that fucosylation of N1 EGF12 may be more sensitive to disruption, the ligand binding defects in *POFUT1 S162L* patient cells may be predominantly from loss of N1 EGF12 *O*-fucosylation rather than effects from N3.

N1 signaling plays an important role in human fibroblasts during tissue repair and fibrosis. Activation of N1 signaling in fibroblasts can promote their proliferation and regulate the expression of several genes involved in fibroblast survival (Ribeiro da Silva, A., Neri, E.A., et al. 2020). N1 also participates in fibroblast differentiation, guiding them to mature into specialized fibroblast subtypes (Wei, K., Korsunsky, I., et al. 2020), or fibroblast-derived cells such as myofibroblasts, in response to appropriate environmental cues mediated by collagen secretion (Dees, C., Tomcik, M., et al. 2011). Furthermore, N1 abnormalities in fibroblasts have been implicated in the pathogenesis and progression of fibroblast-related disorders (Hu, B. and Phan, S.H. 2016). Excessive N1 signaling has been associated with abnormal fibroblast proliferation and fibrosis promotion (Hu, B. and Phan, S.H. 2016). For instance, in fibroblast-derived diseases such as pulmonary fibrosis or renal fibrosis, dysregulated N1 signaling may contribute to the development of fibrotic tissue. Consequently, the ventricular symptoms of the *POFUT1 S162L* patient (Takeuchi, H., Wong, D., et al. 2018), may be caused by the loss of *O*-fucose modification on N1 EGF12.

Endogenous N1 was purified from human primary cells and the levels of glycan modification were analyzed by mass spec. A deficiency of *O*-fucosylation on EGF12 was observed in the *POFUT1S162L* patient fibroblasts. In previous studies, N1 *EGF12f* mice and flies have been described (Ge, C. and Stanley, P. 2008, Pandey, A., Harvey, B.M., et al. 2019). The *EGF12f* fly model demonstrated embryonic lethality accompanied by a distinct loss of Notch phenotype (Harvey, B.M., Rana, N.A., et al. 2016, Pandey, A., Harvey, B.M., et al. 2019). Similarly, *EGF12f* mice with a C57BL/6J background exhibited lethality; however, this lethality was successfully rescued by crossing with mice of a different genetic background (Varshney, S., Wei, H.X., et al. 2019). These findings may provide insight into the survival of the *POFUT1 S162L* proband. Despite the absence of EGF12 *O*-fucose modification in their fibroblast cells, there may be compensatory mechanisms or genetic factors in place that allow for survival even in the absence of these specific modifications.

## Materials and methods

### Cell culture

HEK293T cells were purchased from ATCC. HEK293T cells were cultured in Dulbecco’s Modified Eagle’s Medium (DMEM) with high glucose and 10% bovine calf serum at 37 °C in a humidified incubator at 5% CO_2_. Normal (Control No. 1: GM0038 and Control No. 2: GM3348) and *POFUT1-S162L* fibroblasts (see reference (Takeuchi, H., Wong, D., et al. 2018) for details) were cultured identically to HEK293T cells with 10% fetal bovine serum substituted for bovine calf serum.

### Western blotting analysis

Western blotting methods were described previously (Matsumoto, K., Kumar, V., et al. 2022). Cells were lysed with a solution of 1% NP40 in TBS (Tris-buffered saline, pH 7.4). Lysates were then boiled with SDS-PAGE sample buffer containing 2-mercaptoethanol. Samples were electrophoresed via SDS-PAGE using a 7.5 % pre-cast stain-free gel (Bio-Rad). Protein was transferred to a nitrocellulose membrane using a transfer buffer containing 10% methanol at 100 V for 60 min at room temperature. The membrane was blocked with 5% BSA at room temperature for one hour. Primary antibody staining was performed overnight at 4 °C using the following antibodies and dilutions: POFUT1 (1/2500, #14929-1-AP, Proteintech), human N1 ICD (1/1000, #D1E11, Cell Signaling), Tubulin (1/5000, #DM1A, Sigma), human N1 ECD (1/1000, #AF5317, R and D).

Membrane was washed three times for 5 min with TBS containing 0.1% Tween 20. Secondary antibody staining was performed at room temperature for one hour using a 1/5000 dilution of the antibody (anti-mouse IgG IRDye 680 (1/5000, Li-COR #925-68970), anti-sheep IgG IRDye 800 (1/5000, Themo #SA5-10060)). Membrane was washed three times for 5 min with TBS containing 0.1% Tween 20, followed by a final wash with TBS for 5 min. The protein bands were detected using an Odyssey CLx Imaging System (LI-COR Biosciences). The intensity of the bands was calculated using Image Studio software (LI-COR Biosciences). The ratios of the intensities of each band were calculated, and bar graphs were generated using Excel.

### Ligand binding assay

Ligand binding assays were described previously (Kakuda, S. and Haltiwanger, R.S. 2017, Matsumoto, K., Kumar, V., et al. 2022). Fibroblasts were dissociated using cell dissociation buffer (GIBCO; 13151014) and collected by centrifugation at 3000 × *g*, 4 °C, 5 min. A total of 1x10^5^ cells were washed three times with Hanks’ balanced salt solution containing 1% BSA, 1 mM CaCl_2_, and 0.02% azide (HBSS). For each assay, 1x10^5^ cells were used. Washed cells were incubated with mDLL1-Fc (0-150 nM) and mJAG1-Fc (0-37.5 nM) in HBSS on ice for 1 h. After three washes with HBSS, the Fc portions were incubated with PE-conjugated anti-human Fc (R and D; diluted 1/20) on ice for 1 h. The cells were centrifuged, washed three times with HBSS, and the cell surface PE was measured using the Accuri6 flow cytometer system (BD Biosciences). The intensity of PE for each ligand minus negative control was used for the generation of bar graphs, and each experiment was performed as biological triplicates. Bar graphs were generated in Excel.

### N1 immunopurification from HEK293T cells and fibroblasts

N1 purification was performed essentially as described previously (Matsumoto, K., Kumar, V., et al. 2022). To purify N1, twenty 10 cm dishes of fibroblasts or one 10 cm dish of HEK293T cells were used. Fibroblasts and HEK293T cells were washed three times with TBS and lysed using 1 ml of 1% NP40-TBS buffer supplemented with an EDTA-free cOmplete protease inhibitor tablet (Thermo) per dish. The lysates were incubated on ice for 20 min and then centrifuged at 16,260 x *g* for 5 min at 4 °C to clear the lysate. The resulting supernatant was used for immunopurification (IP) of endogenous human N1. For IP, 15 μg of human N1 antibody (AF5317: R&D Systems) was covalently coupled to 60 μl of Protein G-Dynabeads (Thermo Fisher Scientific) using a BS3 cross-linker. The coupled beads, bound to 5 μg of anti-N1, were incubated with the cell lysate for 4 to 6 hours in a cold room with tilting rotation. After IP, the Dynabeads were washed three times with cold 1% NP40-TBS and three times with cold TBS. To elute N1, the Dynabeads were incubated with 8M urea at 37 °C for 5 min.

### Glycoproteomic mass spectral analysis of Human N1

The reduction, alkylation, digestion, and mass spectral analyses were performed following our previously established protocol (Kakuda, S. and Haltiwanger, R.S. 2017, Matsumoto, K., Kumar, V., et al. 2022). Immunopurified human N1 was reduced with 25 mM TCEP (Thermo) in 8 M urea and heated at 100 °C for 5 min. After cooling to room temperature, iodoacetamide was added to 25 mM and the sample was incubated in the dark for 30 min. The sample was then diluted 8-fold using mass spectral grade water. Diammonium phosphate was added to a final concentration of 20 mM from a 1 M stock, and the sample was split into three fractions. Fractions were digested at 37 °C for 4-6 hours with 500 ng of trypsin (Sigma), chymotrypsin (Thermo), or V8 (Thermo) protease. Peptides were desalted using Pierce C18 Spin Tips (Thermo), washed with 0.1% formic acid, and eluted with 50% acetonitrile in 0.1% formic acid. Peptide separation was carried out using an Easy nano-LC HPLC system with a C18 EasySpray PepMap RSLC C18 column (50 mm × 15 cm, Thermo). The separation of glycopeptides was performed using a 30-minute binary gradient consisting of solvent A (0.1% formic acid in water) and solvent B (90% acetonitrile and 0.1% formic acid in water) at a constant flow rate of 300 nL/min. Peptides were detected using a Q Exactive Plus mass spectrometer (Thermo Fisher Scientific). Higher energy collisional dissociation-tandem mass spectrometry (HCD-MS/MS) was employed, where the 10 most abundant precursor ions in each MS scan were selected for fragmentation (collision energy set at 27%, 2 × 10^5^ gain control, isolation window m/z 3.0, dynamic exclusion enabled, and 17,500 fragment resolution). Peak lists were generated using Xcalibur software with default settings. RAW data files were analyzed using Proteome Discoverer 2.1.0.81 (Thermo) and searched against a human N1 ECD database (Q01705, 18 April 2012—v3). Byonic software version 2.10.5 (Protein Metrics) was incorporated as a node inside Proteome Discoverer for identifying peptides with glycan modifications. Peptide spectral matches (PSMs) with a Byonic score below 200 were culled prior to analysis for the data dependent search analysis. The search allowed for two missed cleavages, with a fixed modification set as carbamidomethyl (+57.021464) on cysteines and variable modifications including oxidation (+15.994915) on methionine, histidine, asparagine, or aspartic acid. Glycoforms were searched as rare-2 modifications, including dHex (+146.057909), HexNAc-dHex (+349.137281), Hex-HexNAc-dHex (+511.190105), NeuAc-Hex-HexNAc-dHex (+802.285522), Hex (+162.052824), Pent-Hex (+294.095082), Pent-Pent-Hex (+426.137341), HexNAc (+203.079373), Hex-HexNAc (+365.132196), and NeuAc-Hex-HexNAc (+655.227613).

The glycoform stoichiometry on each EGF repeat was quantified based on the area under the curve of each EIC for all biological and technical replicates. Byonic search results are provided in Tables S1-S3. Table S3 provides results of a targeted search for EGF8 and EGF12 peptides using the following m/z: for EGF8 peptide, unmodified (m/z 1081.07), monosaccharide (m/z 1129.76); for EGF12 peptide, unmodified (m/z 1308.87), monosaccharide (m/z 1357.55), disaccharide (m/z 1069.19), trisaccharide (m/z 1109.7), tetrasaccharide (m/z 1182.47). PSMs with a Byonic score below 150 were culled prior to analysis for the Targeted search analysis.

### Statistics

Data are presented as mean ±standard deviation. Calculations were performed using the unpaired t-test calculated by Excel.

## Data availability

The mass spectrometry proteomics data have been deposited to the ProteomeXchange Consortium via the PRIDE (Perez-Riverol, Y., Bai, J., et al. 2022) partner repository with the data set identifier PXD050256.

## Author contributions

KM and RSH conceived of the project; KM and RSH designed the research; RSH provided funding; KM performed the experiments; KM wrote the original draft; KM, KBL, and RSH interpreted results and edited the paper.

## Funding and additional information

Supported by funding from NIH (R01 GM061126 and R35 GM148433 to RSH) and the Georgia Research Alliance.

## Acknowledgments

Dr. Hudson Freeze for reagents and helpful discussions, Drs. Pamela Stanley, Hamed Jafar-Nejad, Kazuhiro Aoki, Michael Tiemeyer, and Mayumi Ishihara for helpful discussions.

