## Supplement text for "Analysis of endogenous NOTCH1 from *POFUT1 S162L* patient fibroblasts reveals the importance of the *O*-fucose modification on EGF12 in human development"

Contents:

Supplemental Tables S1-S4

Supplemental Figures S1-S4

**Supplemental Table legends:**

**Table S1. Mass spectral analysis of peptides resulting from tryptic, chymotryptic, or V8 digestion of endogenous NOTCH1 from HEK293T cells**

**Table S2. Mass spectral analysis of NOTCH1 chymotryptic peptides from normal or *POFUT1 S162L* patient fibroblasts (data dependent search)**

**Table S3. Mass spectral analysis of NOTCH1 chymotryptic peptides from normal or *POFUT1 S162L* patient fibroblasts (Targeted search for EGF8 and 12 peptides)**

**Table S4: Description of the files uploaded to the PRIDE repository.**

### Supplemental Figure legends:

**Figure S1. Anti-hN1 ECD antibody can immunopurify human N1.**

Endogenous human N1 was immunopurified from HEK293T cells (**A**), control fibroblasts (**B**), and *POFUT1 S162L* fibroblasts (**C**) with anti-human N1 ECD antibody as described in Materials and Methods. Western blot with anti-N1 ECD of Input, After IP, Wash, and Elution is shown. Molecular weight markers are shown on the left in kD.

**Figure S2. EICs showing relative levels of *O*-fucose glycoforms on peptides from N1 EGF repeats isolated from fibroblasts**.

Each EIC was generated using Xcalibur as described in Materials and Methods. Black (unmodified), red (monosaccharide), blue (disaccharide), green (trisaccharide), and purple (tetrasaccharide) lines indicate the curves for the various glycoforms. Mass spectral data can be found in Fig. S4 and Table S2.

**Figure S3. Annotated MS/MS spectra for all *O*-fucosylated peptides from N1 isolated from HEK293T cells**

N1 was immunopurified from HEK293T cells, digested, and analyzed by nano-LC-MS/MS as described in Experimental Procedures. Glycopeptides were identified using

Byonic. Due to the lability of the fucose-peptide bond in HCD experiments, Byonic is often unable to correctly assign the *O*-fucosylated Ser/Thr residue in a peptide and will often consolidate two separate glycans into a single predicted modification. Peptide fragment ions that have lost the *O*-fucose (a neutral loss) are indicated with a ~. **A**. monosaccharide glycoform of EGF2 peptide. **B**. monosaccharide glycoform of EGF3 peptide. **C**. monosaccharide glycoform of EGF5 peptide. **D**. monosaccharide glycoform of EGF6 peptide. **E**. monosaccharide glycoform of EGF8 peptide with oxidation. **F**. monosaccharide glycoform of EGF9 peptide. **G**. trisaccharide glycoform of EGF12 peptide with *O*-glucose trisaccharide. **H**. disaccharide glycoform of EGF12 peptide with *O*-glucose trisaccharide. **I**. monosaccharide glycoform of EGF12 peptide with *O*-glucose trisaccharide. **J**. tetrasaccharide glycoform of EGF16 peptide. **K**. trisaccharide glycoform of EGF16 peptide. **L.** monosaccharide glycoform of EGF16 peptide. **M**. monosaccharide glycoform of EGF20 peptide. **N**. monosaccharide glycoform of EGF23 peptide. **O**. monosaccharide glycoform of EGF24 peptide. **P.** unmodified EGF24 peptide. **Q**. monosaccharide glycoform of EGF26 peptide. **R**. monosaccharide glycoform of EGF27 peptide with *O*-glucose monosaccharide. **S**. monosaccharide glycoform of EGF34 peptide. **T**. monosaccharide glycoform of EGF35 peptide. **U**. monosaccharide glycoform of EGF36 peptide with an *O*-GlcNAc disaccharide. **V**. monosaccharide glycoform of

EGF36 peptide with an *O*-GlcNAc monosaccharide. N1 EGF36 peptide is consistently modified by *O*-GlcNAc as was shown previously (Kakuda, S. and Haltiwanger, R.S. 2017, Matsumoto, K., Kumar, V., et al. 2022). Green arrows show the oxonium ions from HexNAc from the Fringe modification of *O*-fucose.

**Figure S4. Annotated MS/MS spectra for all *O*-fucosylated peptides from N1 isolated from fibroblast cells**

N1 was immunopurified from control and patient fibroblast cells, digested, and analyzed by nano-LC-MS/MS as described in Experimental Procedures. Glycopeptides were identified using Byonic. Due to the lability of the fucose-peptide bond in HCD experiments, Byonic is often unable to correctly assign the *O*-fucosylated Ser/Thr residue in a peptide. Peptide fragment ions that have lost the *O*-fucose are indicated with a ~. **A**, monosaccharide glycoform of EGF2 peptide from control fibroblast with *O*-glucose trisaccharide. **B**, monosaccharide glycoform of EGF8 peptide from control fibroblast with oxidation. **C**, monosaccharide glycoform of EGF9 peptide from control fibroblast. **D**, unmodified EGF9 peptide from *POFUT1 S162L* patient fibroblast. **E**, trisaccharide glycoform of EGF12 peptide with *O*-glucose trisaccharide from control fibroblast. **F**, unmodified EGF12 peptide with *O*-glucose trisaccharide from *POFUT1 S162L* patient fibroblast. **G**, monosaccharide glycoform of EGF35 peptide from control fibroblast. Green arrows show the oxonium ions from HexNAc from the Fringe modification of *O*-fucose.
