## Supplement Figure for "Analysis of endogenous NOTCH1 from *POFUT1 S162L* patient fibroblasts reveals the importance of the *O*-fucose modification on EGF12 in human development"

### Slide 1
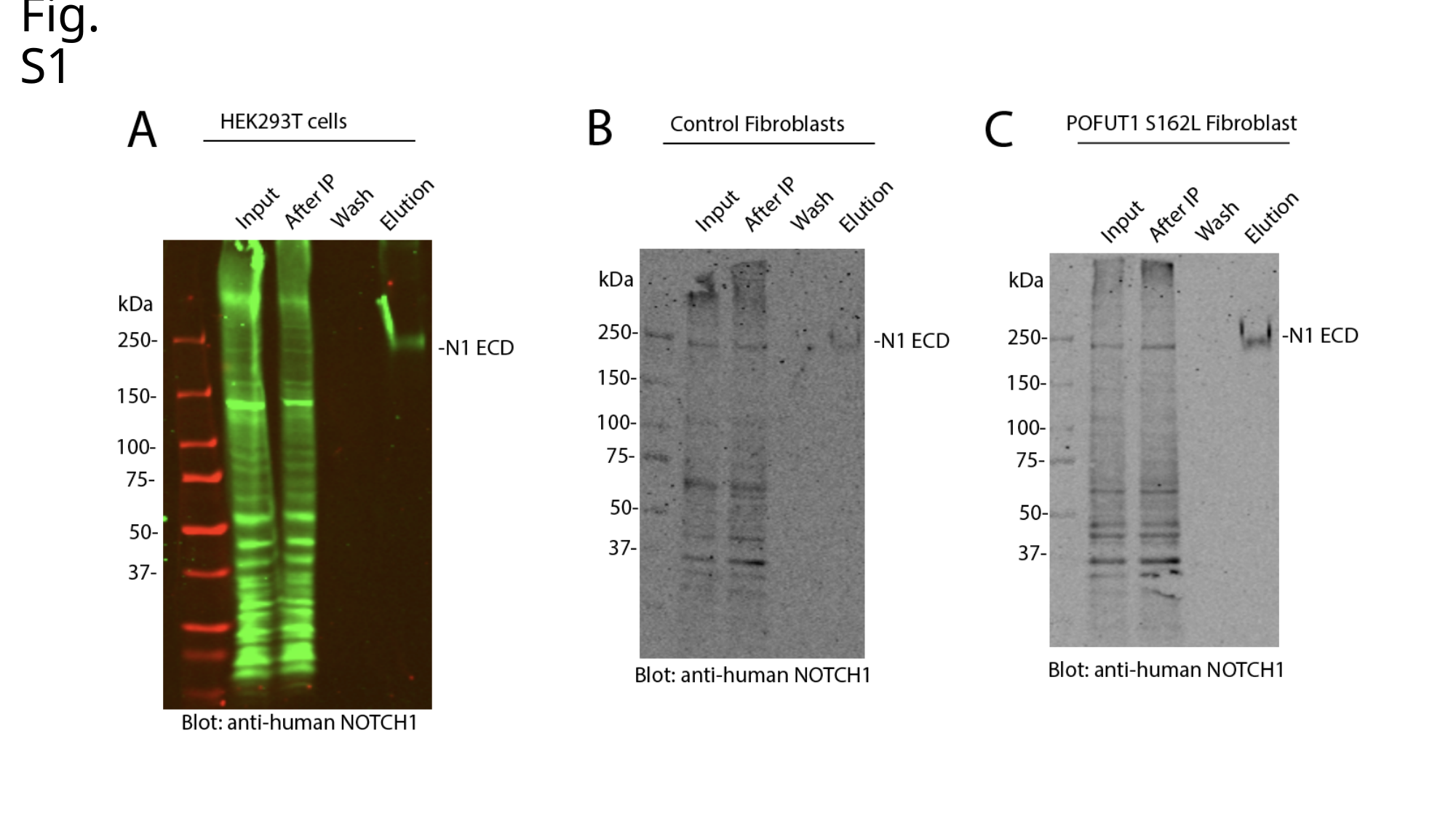

Fig. S1

### Slide 2
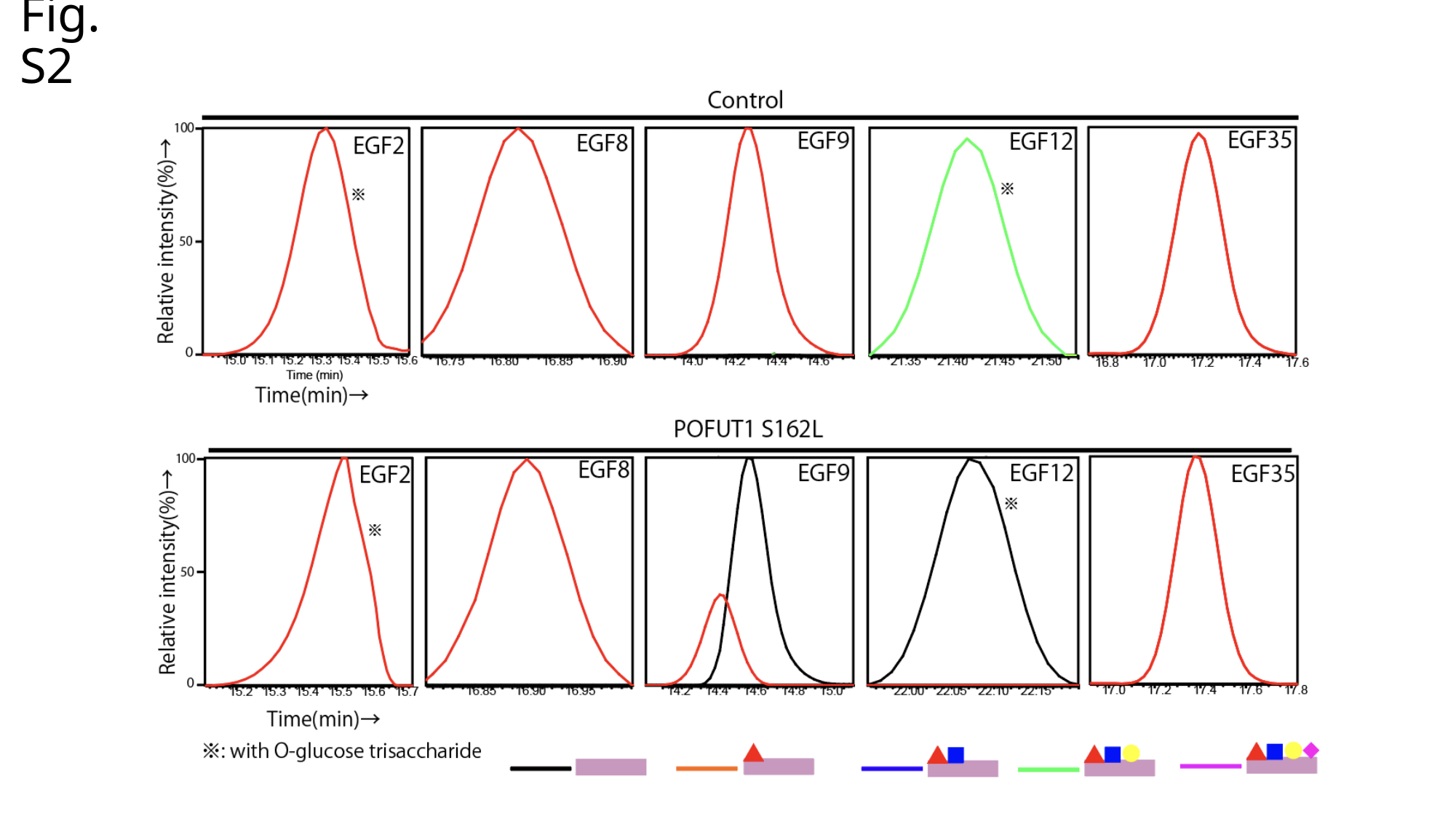

Fig. S2

### Slide 3
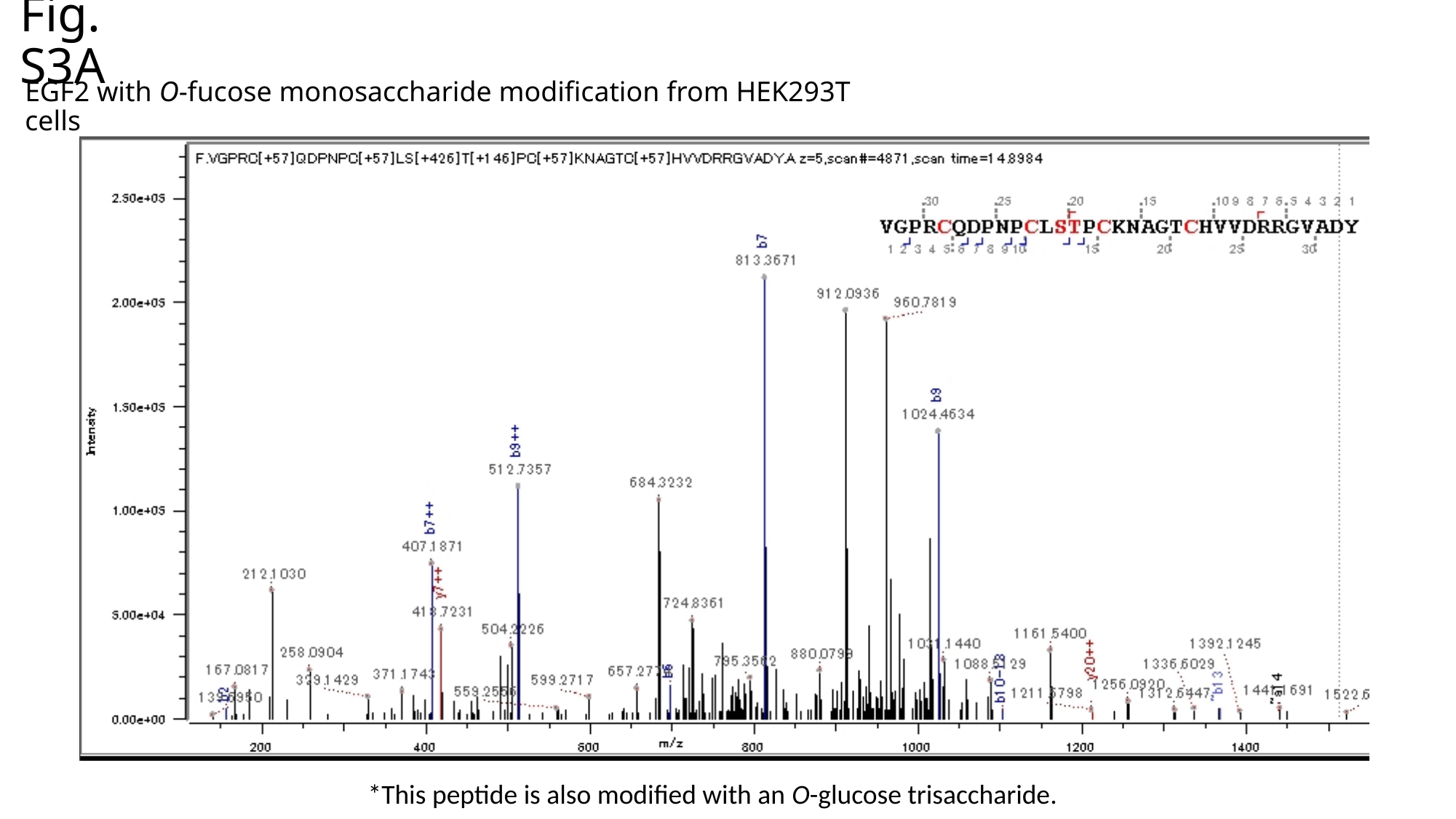

Fig. S3A
EGF2 with O-fucose monosaccharide modification from HEK293T cells
*This peptide is also modified with an O-glucose trisaccharide.

### Slide 4
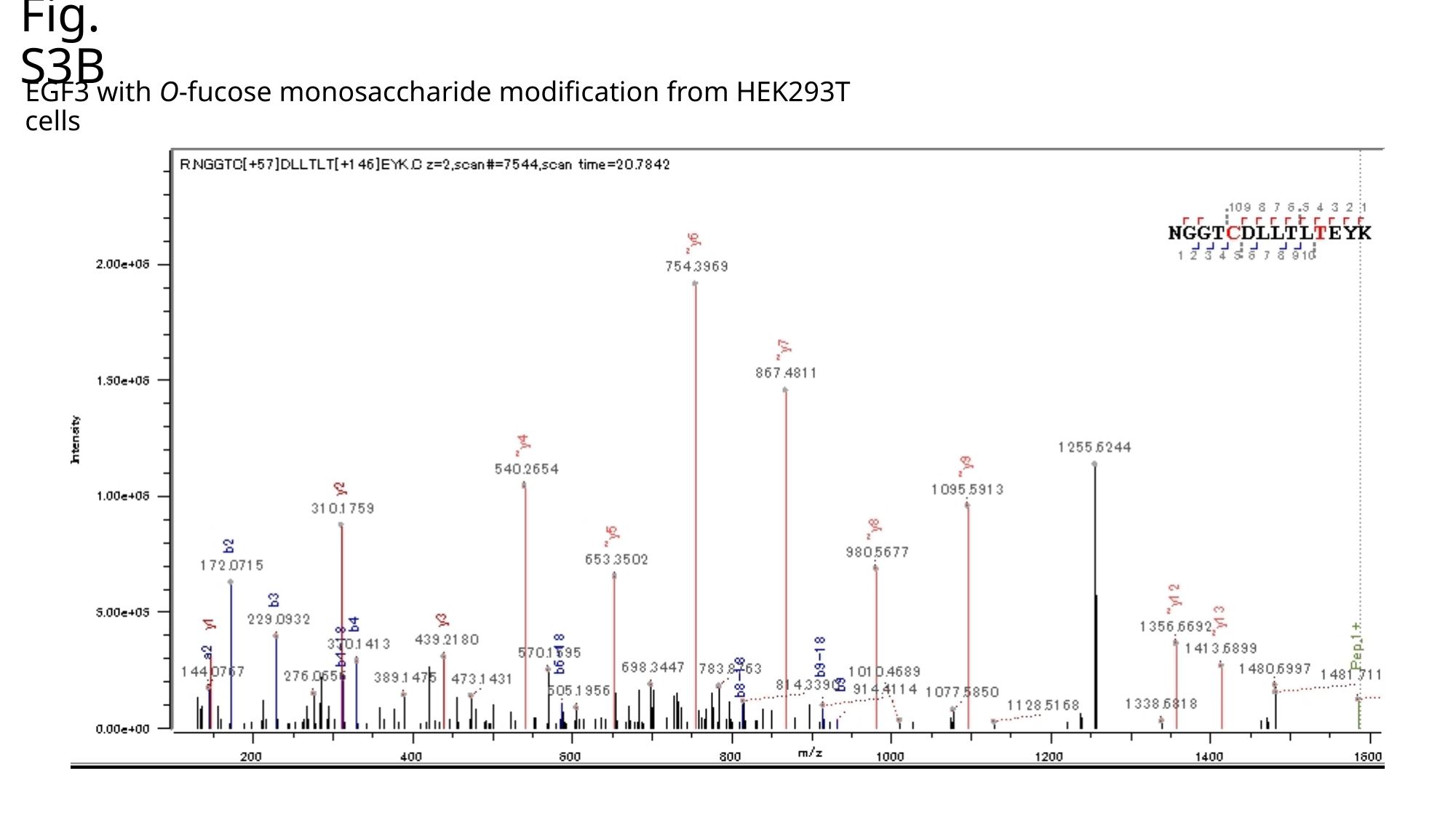

Fig. S3B
EGF3 with O-fucose monosaccharide modification from HEK293T cells

### Slide 5
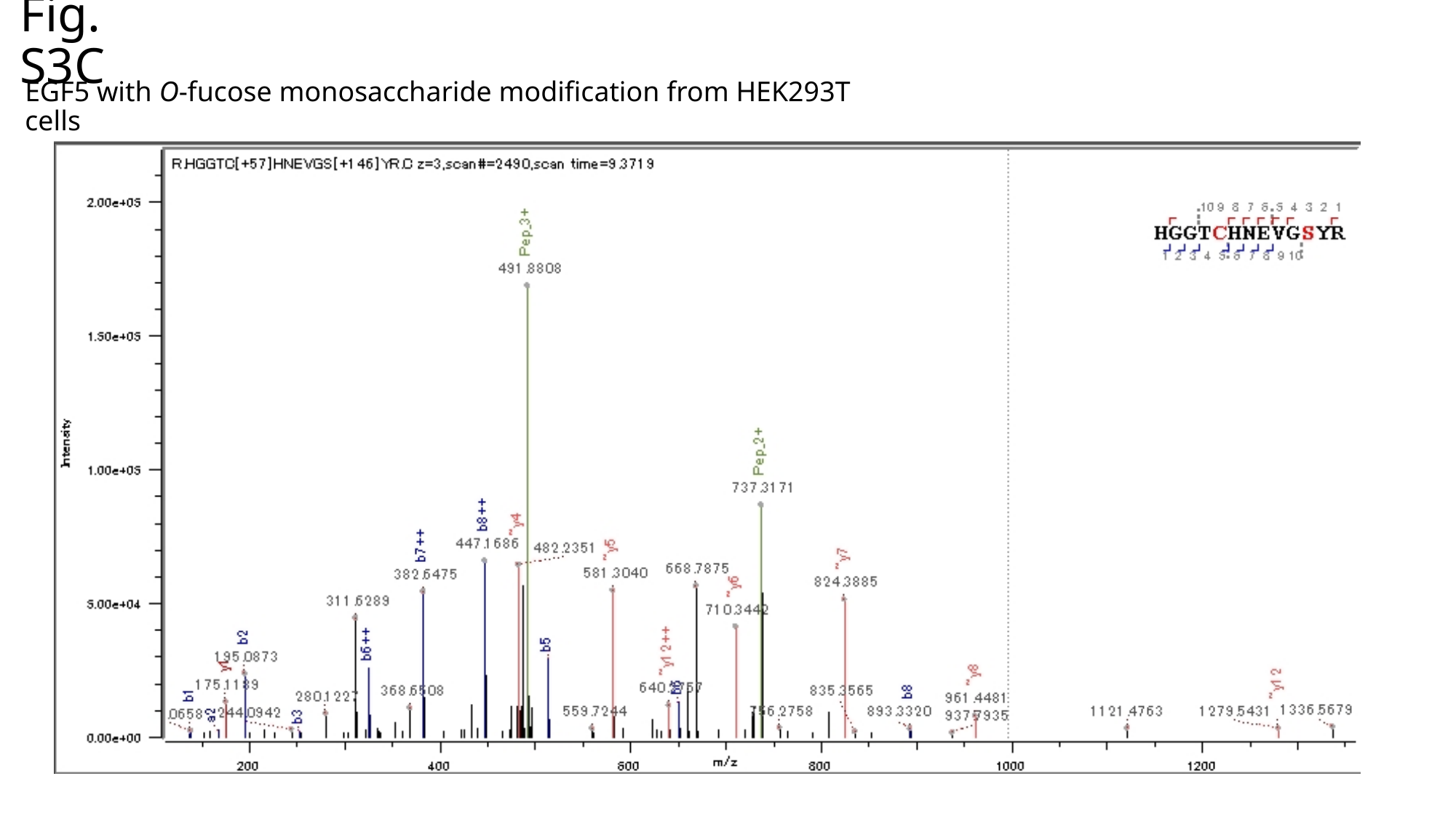

Fig. S3C
EGF5 with O-fucose monosaccharide modification from HEK293T cells

### Slide 6
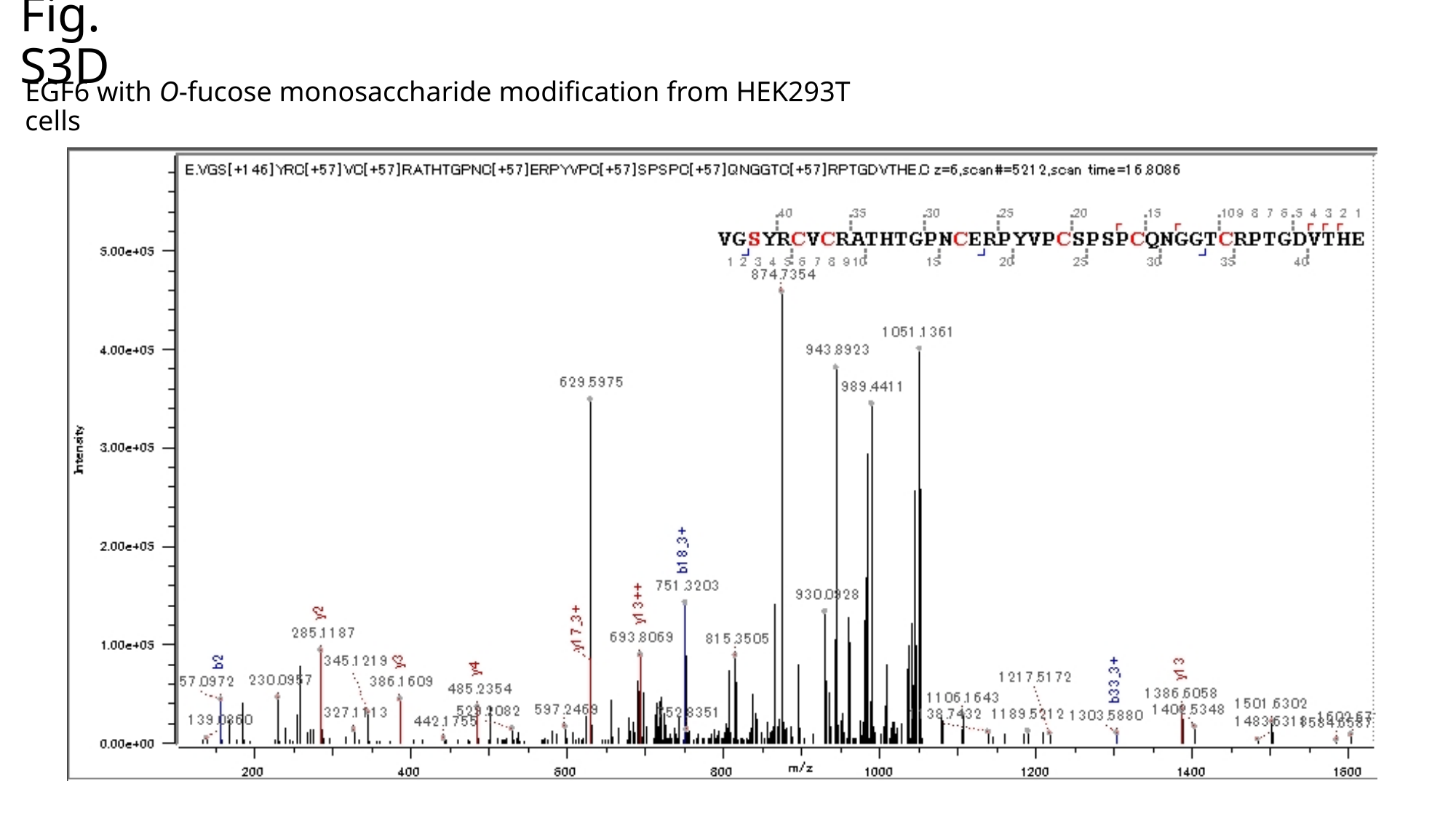

Fig. S3D
EGF6 with O-fucose monosaccharide modification from HEK293T cells

### Slide 7
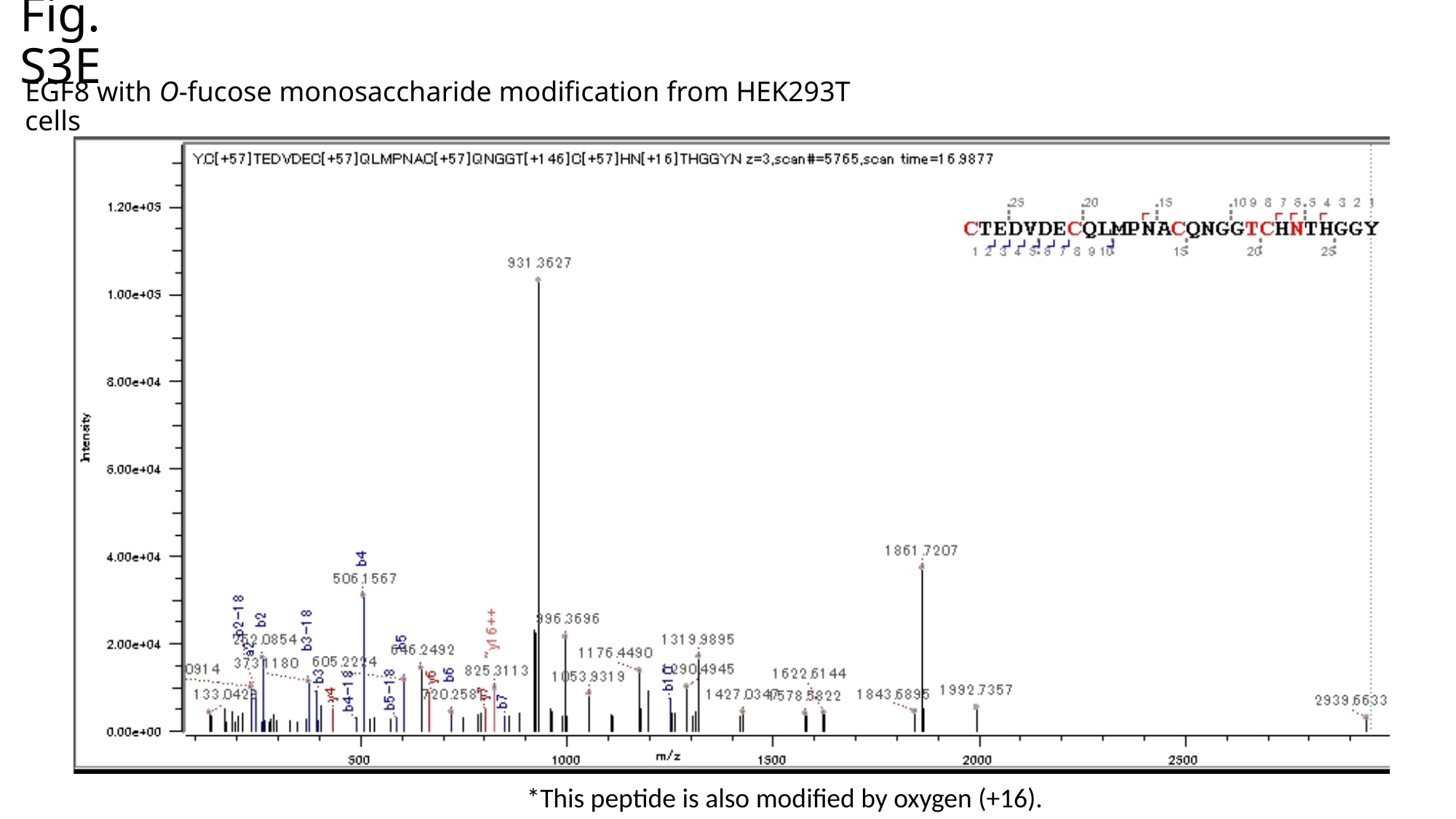

Fig. S3E
EGF8 with O-fucose monosaccharide modification from HEK293T cells
*This peptide is also modified by oxygen (+16).

### Slide 8
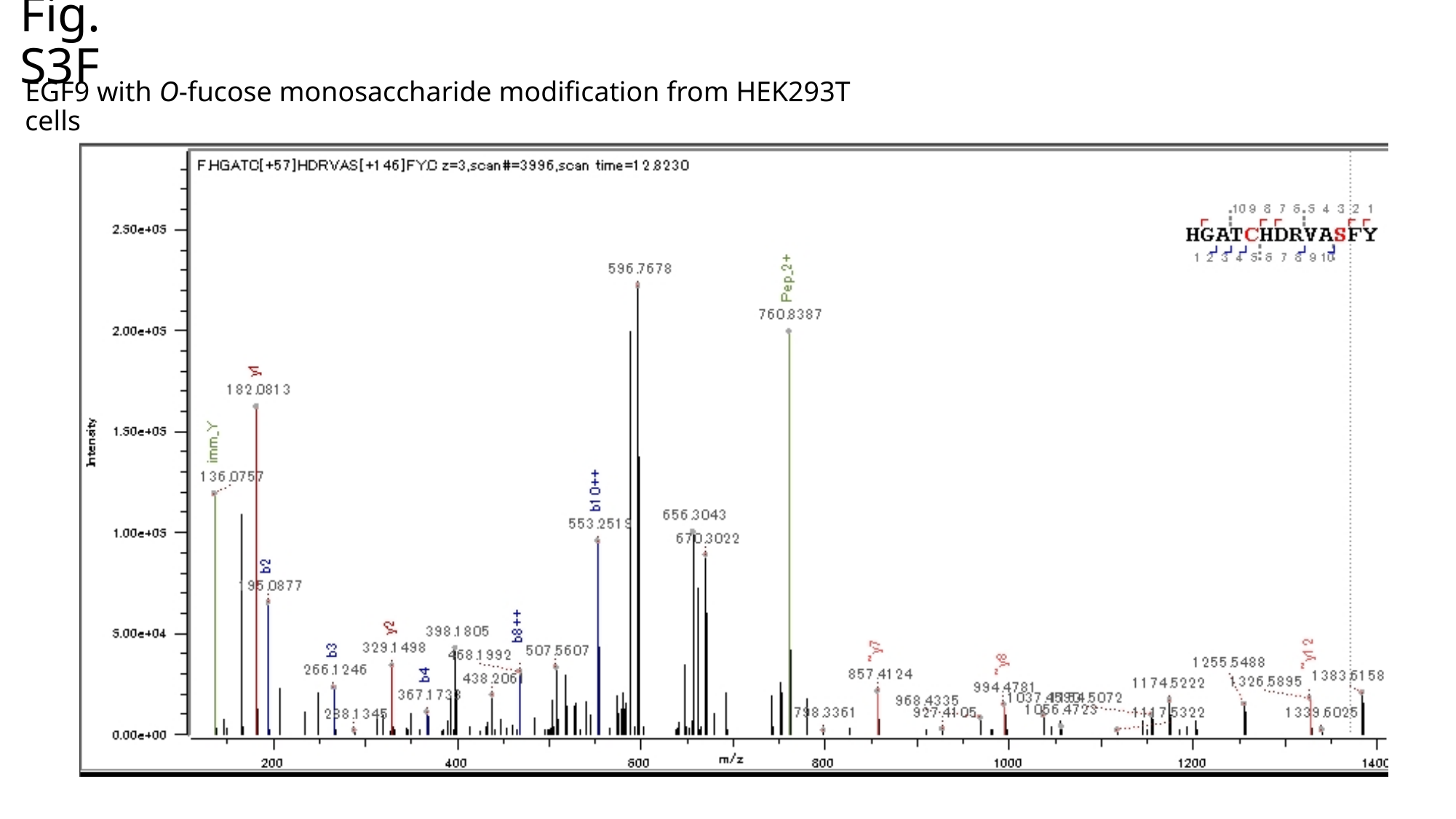

Fig. S3F
EGF9 with O-fucose monosaccharide modification from HEK293T cells

### Slide 9
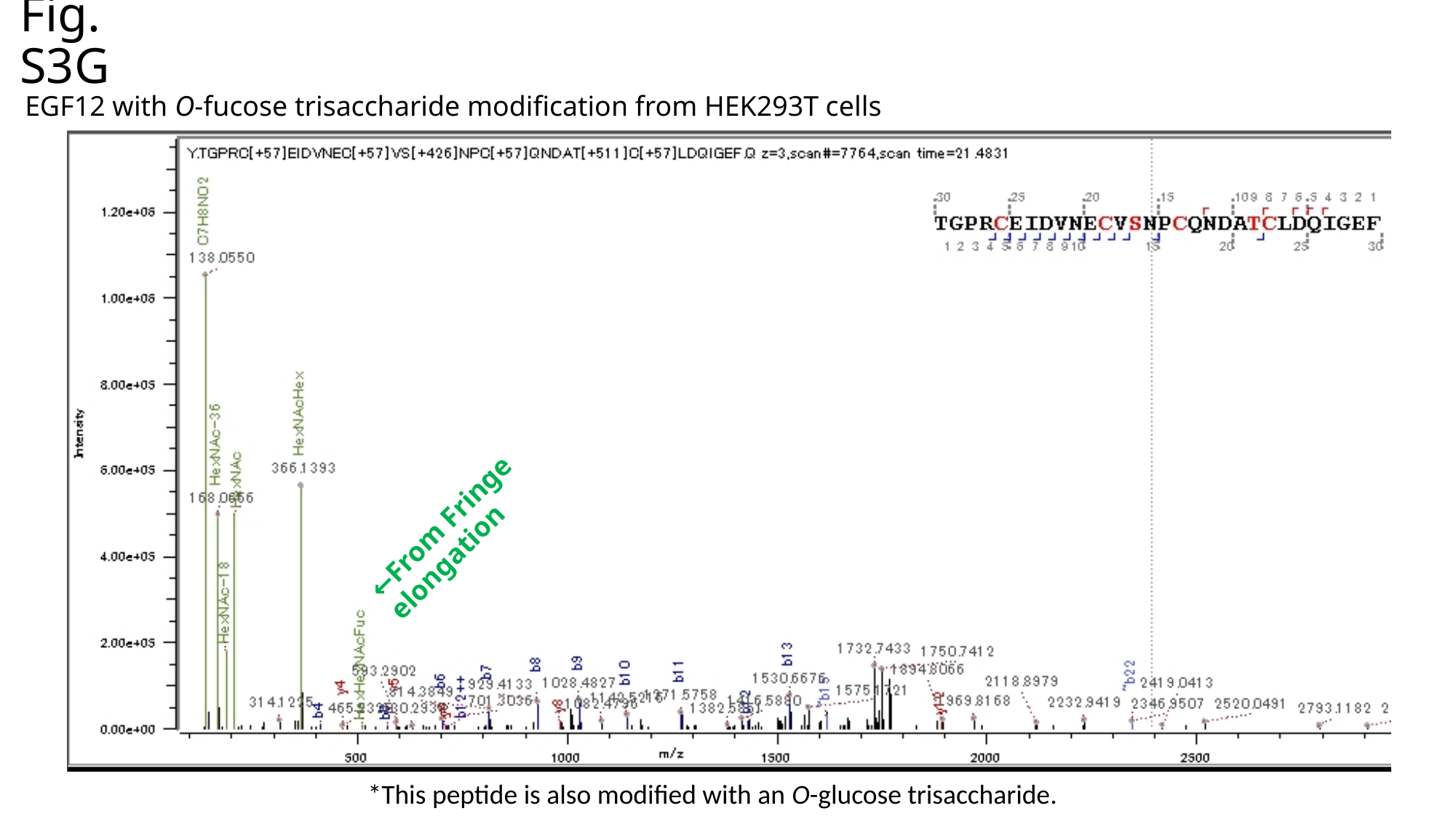

Fig. S3G
EGF12 with O-fucose trisaccharide modification from HEK293T cells
←From Fringe elongation
*This peptide is also modified with an O-glucose trisaccharide.

### Slide 10
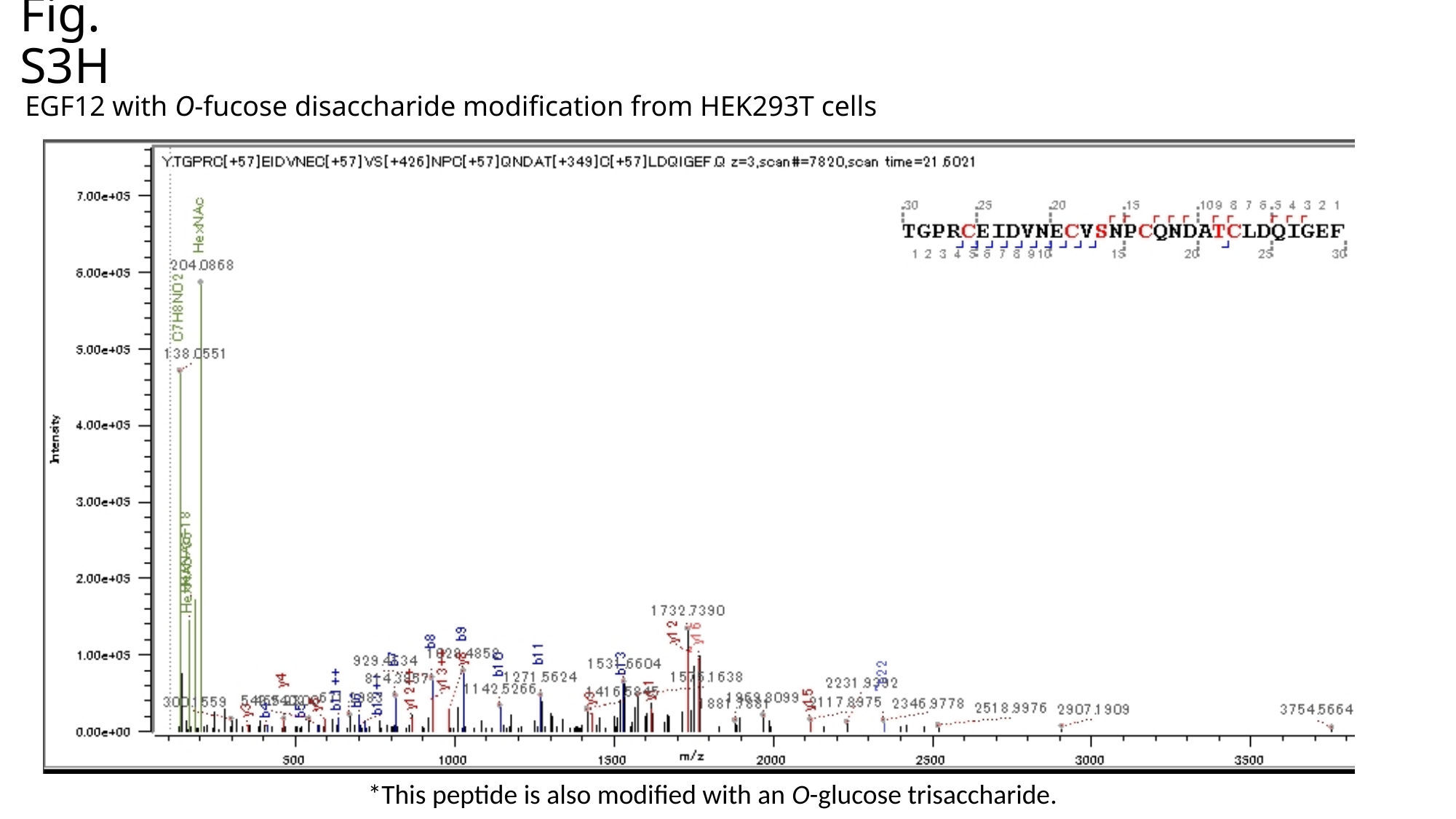

Fig. S3H
EGF12 with O-fucose disaccharide modification from HEK293T cells
*This peptide is also modified with an O-glucose trisaccharide.

### Slide 11
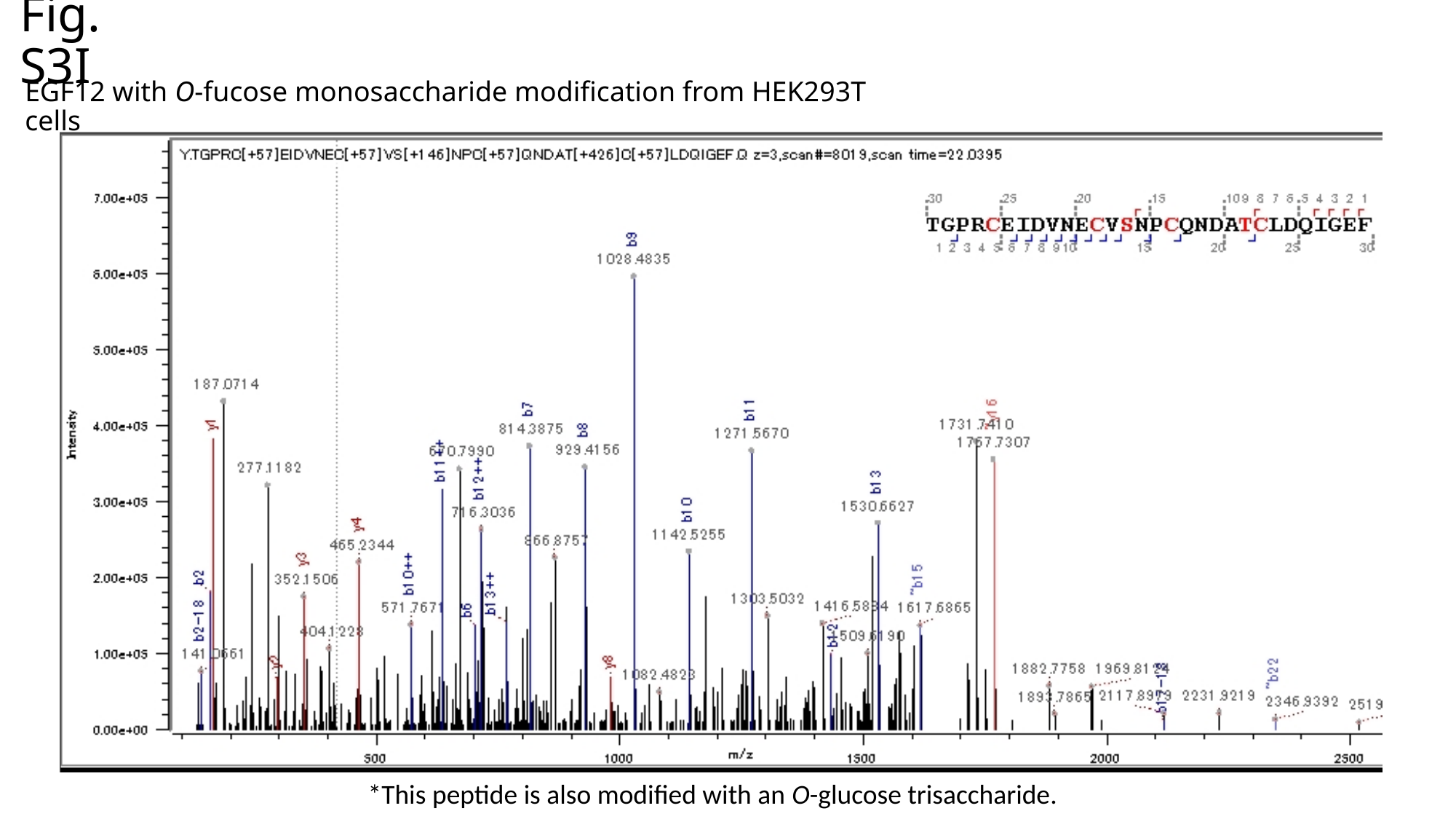

Fig. S3I
EGF12 with O-fucose monosaccharide modification from HEK293T cells
*This peptide is also modified with an O-glucose trisaccharide.

### Slide 12
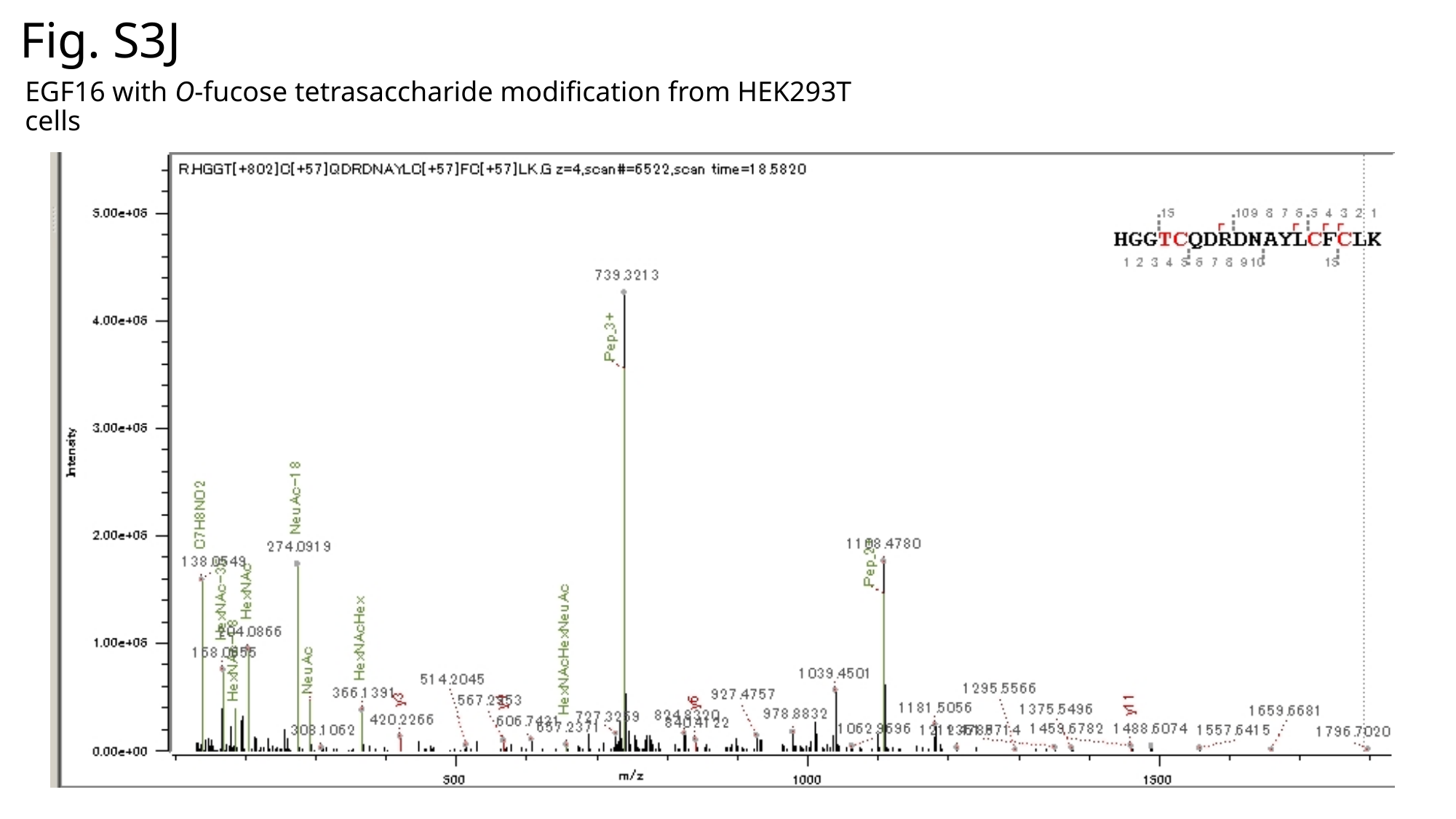

Fig. S3J
EGF16 with O-fucose tetrasaccharide modification from HEK293T cells

### Slide 13
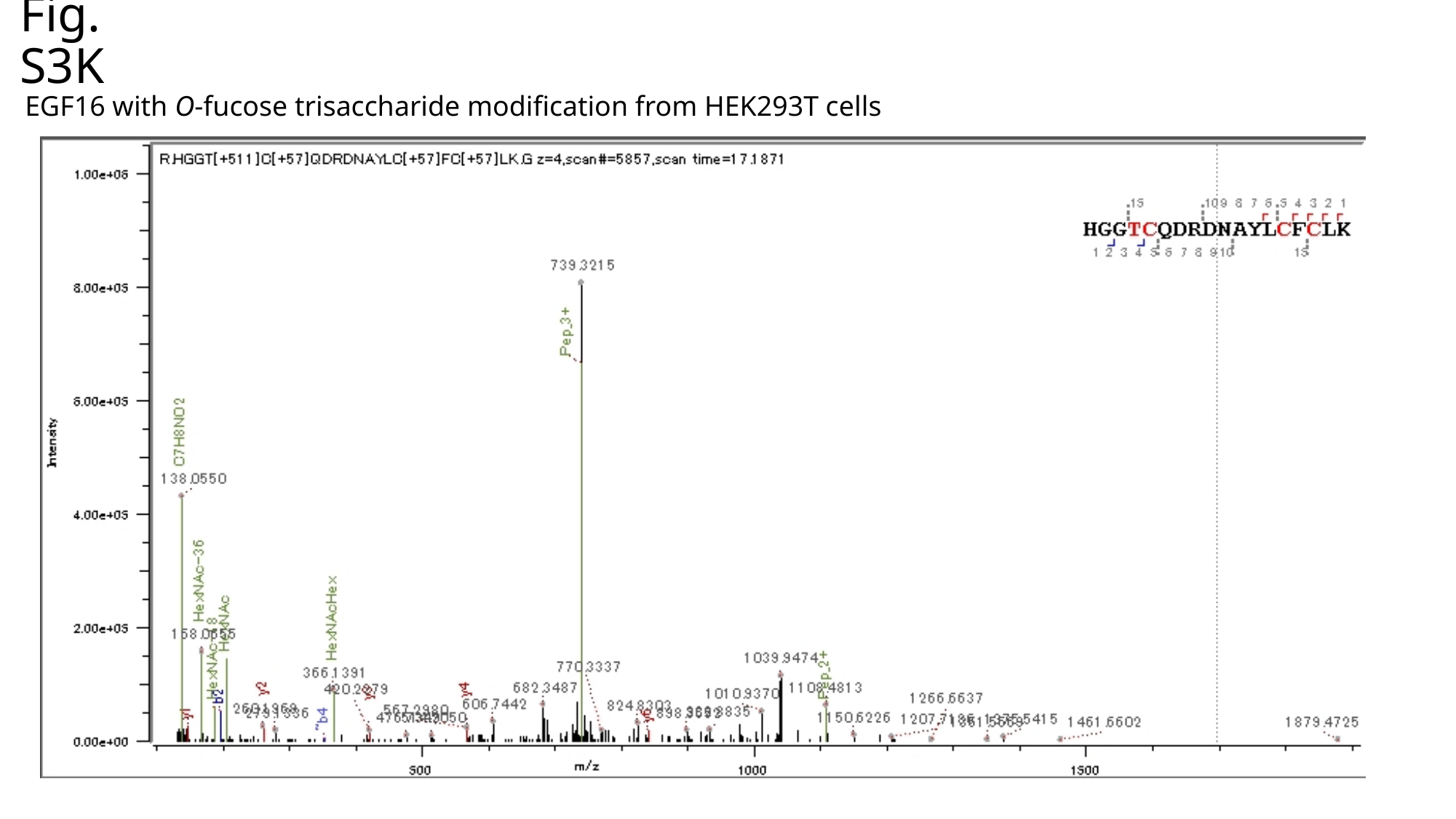

Fig. S3K
EGF16 with O-fucose trisaccharide modification from HEK293T cells

### Slide 14
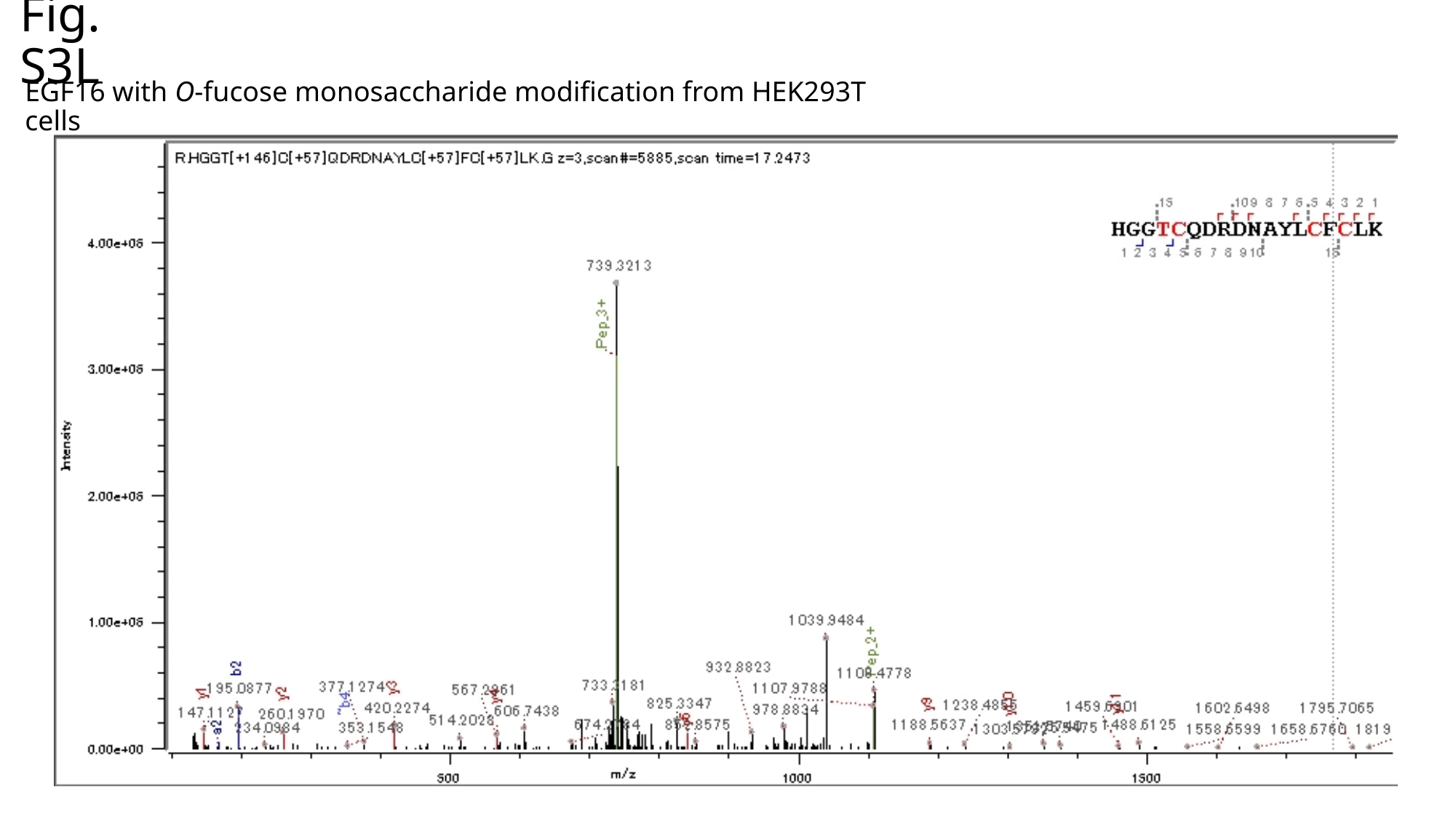

Fig. S3L
EGF16 with O-fucose monosaccharide modification from HEK293T cells

### Slide 15
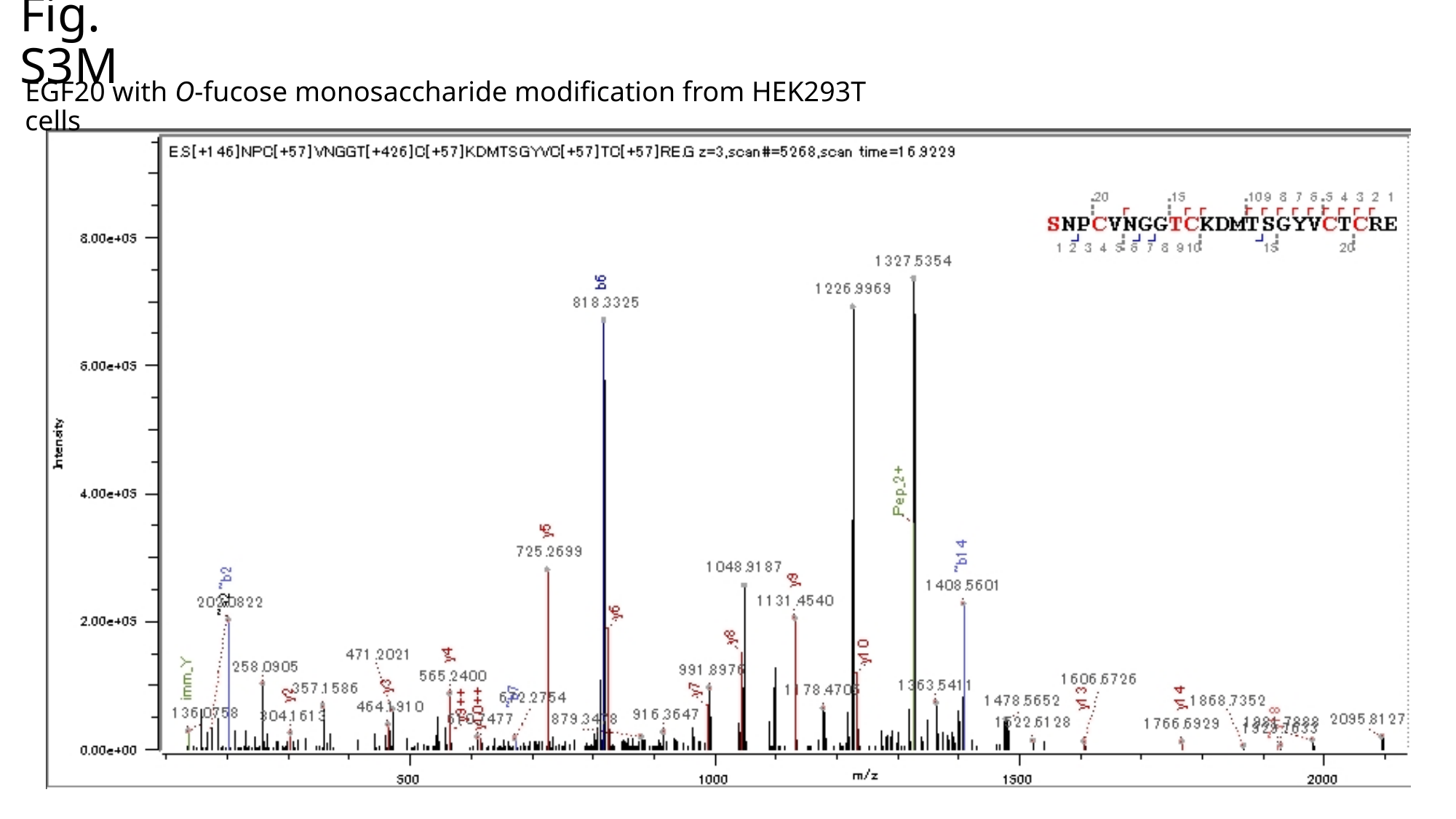

Fig. S3M
EGF20 with O-fucose monosaccharide modification from HEK293T cells

### Slide 16
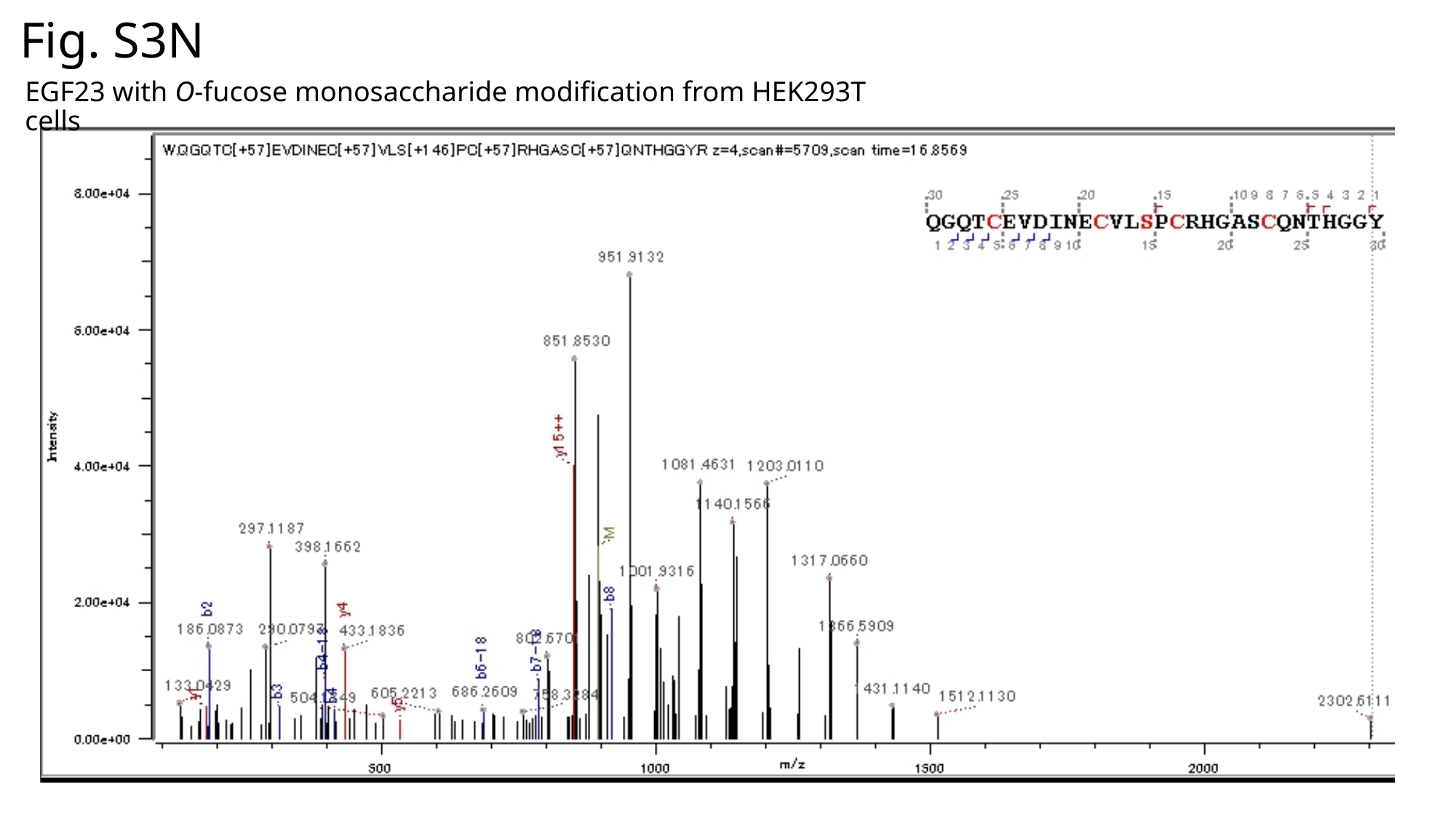

Fig. S3N
EGF23 with O-fucose monosaccharide modification from HEK293T cells

### Slide 17
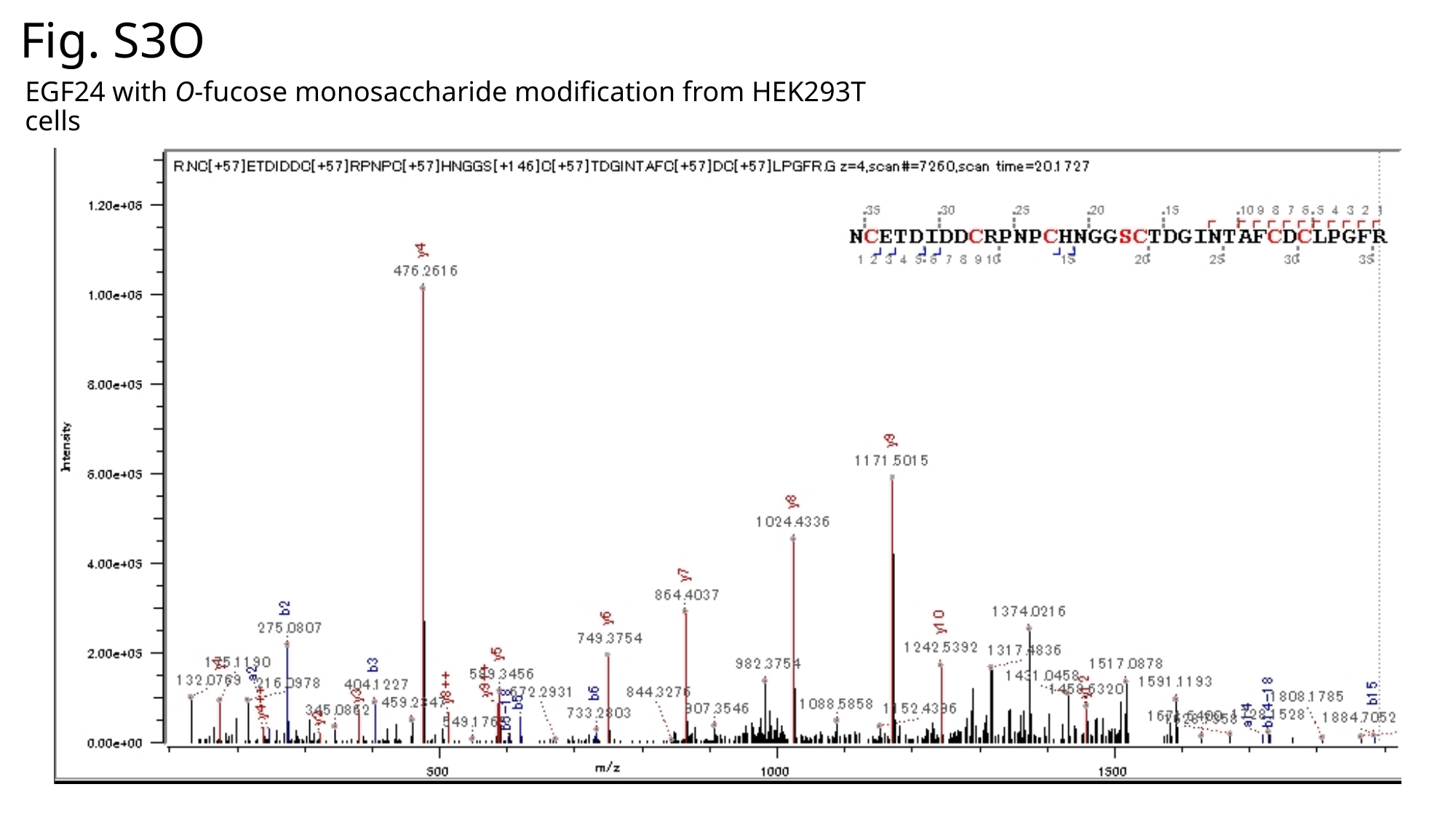

Fig. S3O
EGF24 with O-fucose monosaccharide modification from HEK293T cells

### Slide 18
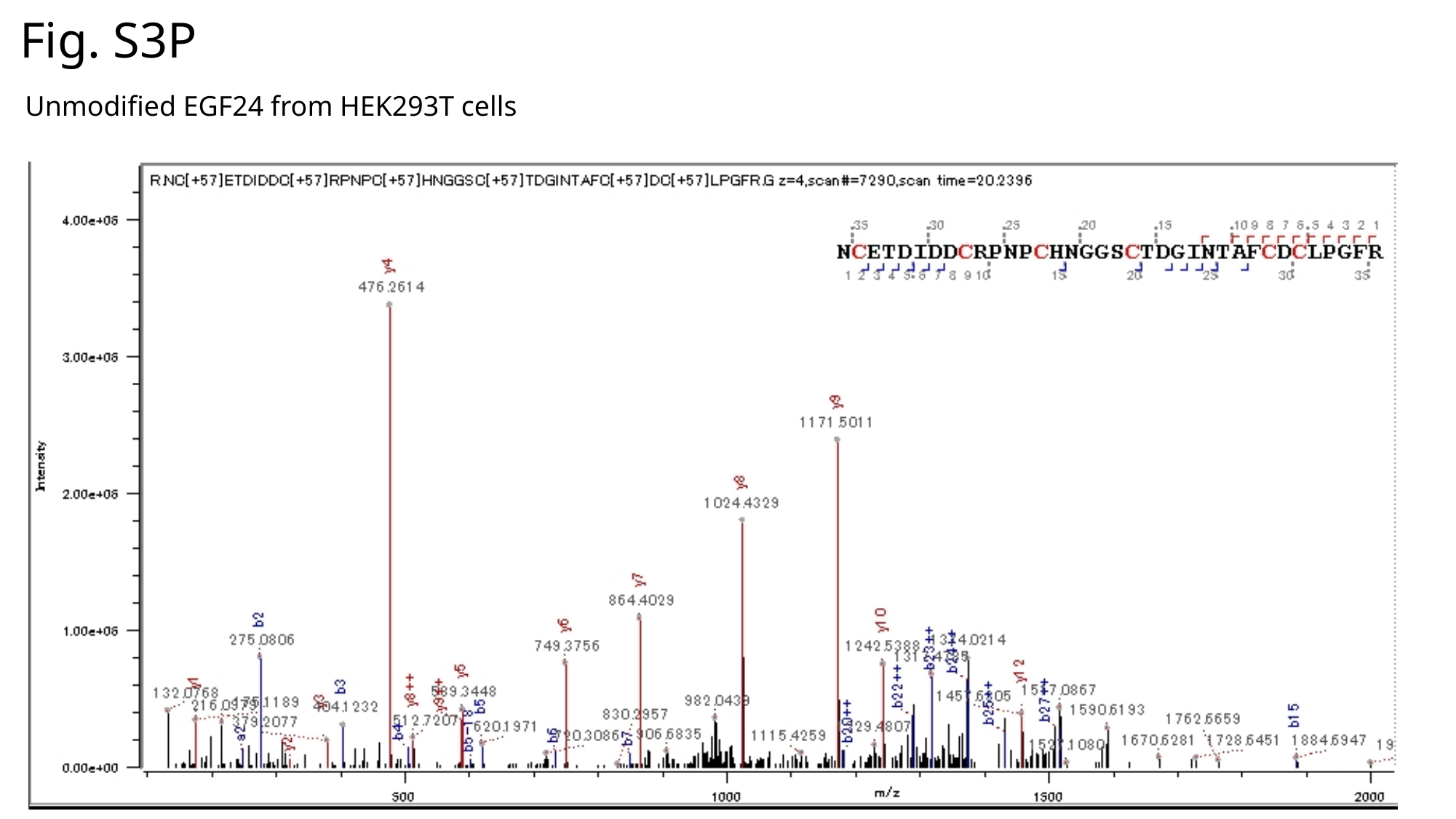

Fig. S3P
Unmodified EGF24 from HEK293T cells

### Slide 19
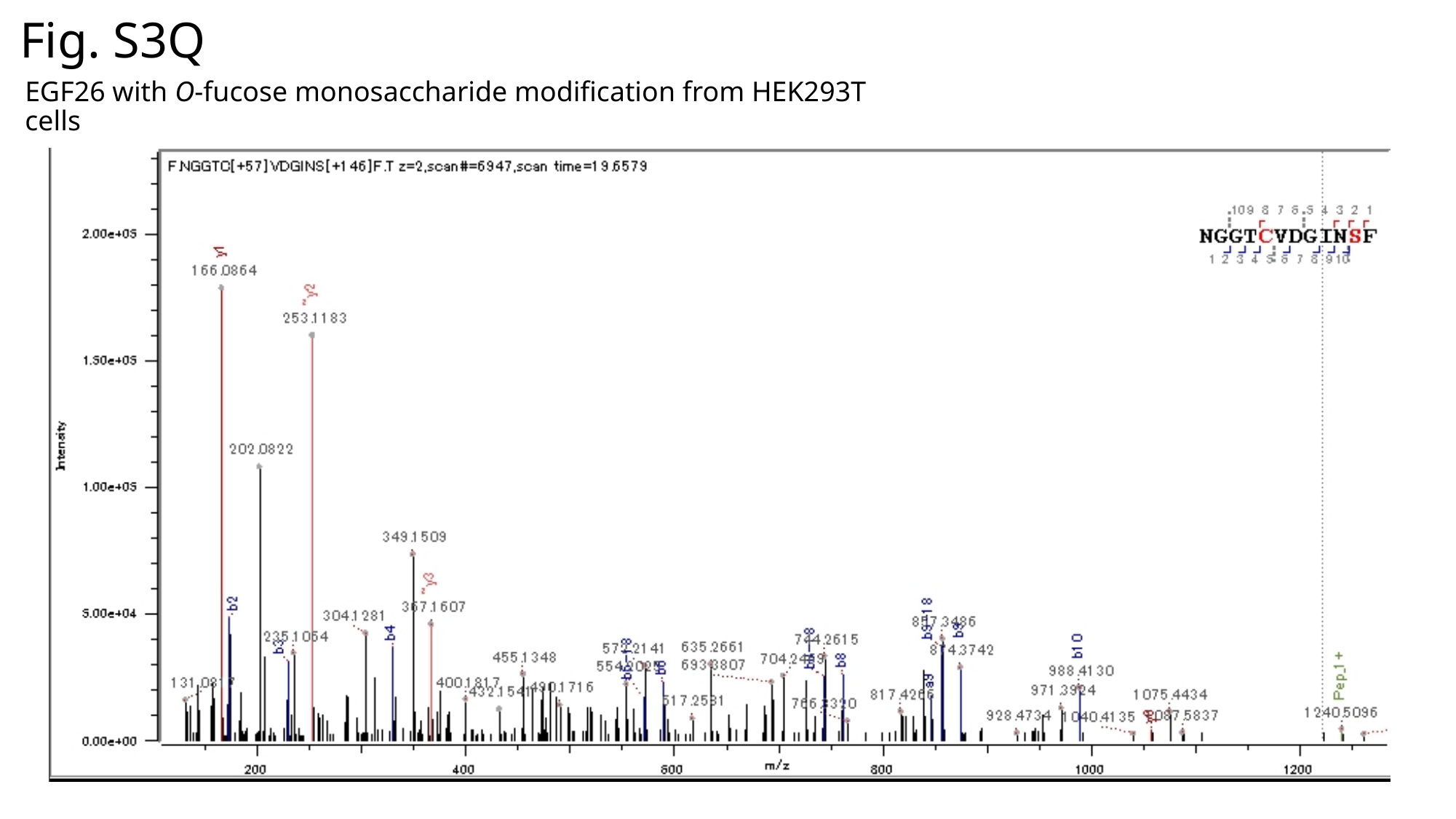

Fig. S3Q
EGF26 with O-fucose monosaccharide modification from HEK293T cells

### Slide 20
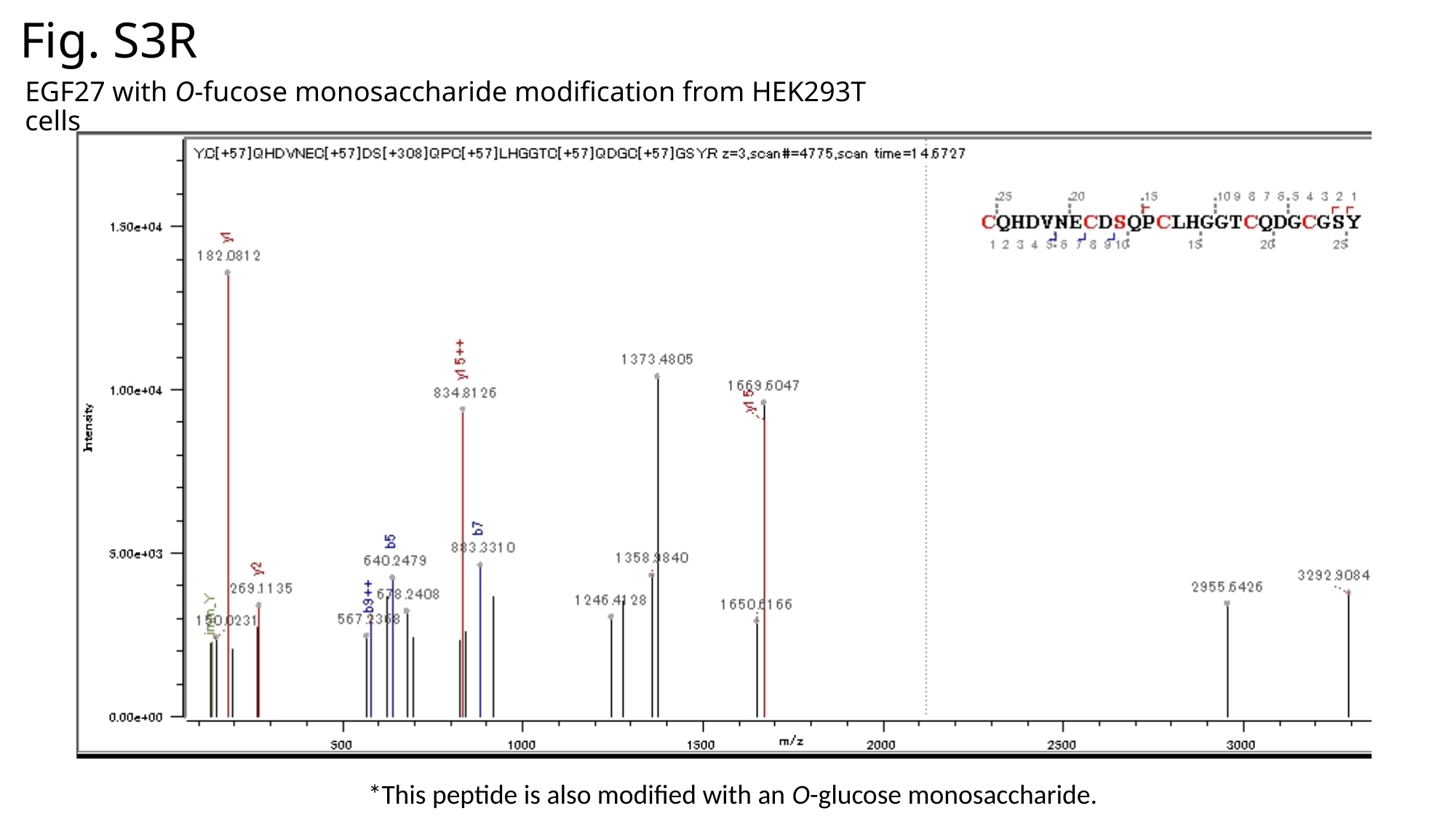

Fig. S3R
EGF27 with O-fucose monosaccharide modification from HEK293T cells
*This peptide is also modified with an O-glucose monosaccharide.

### Slide 21
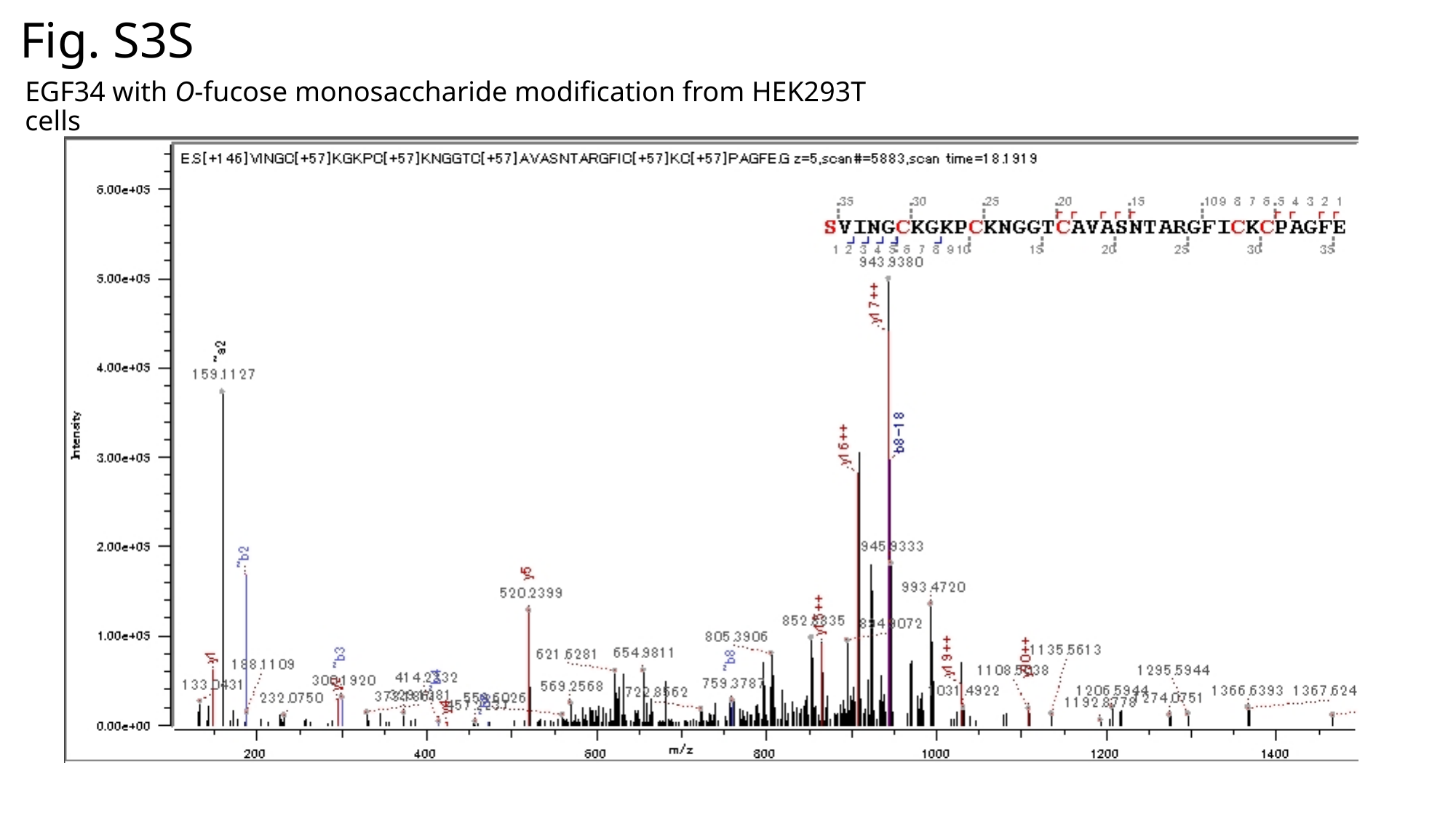

Fig. S3S
EGF34 with O-fucose monosaccharide modification from HEK293T cells

### Slide 22
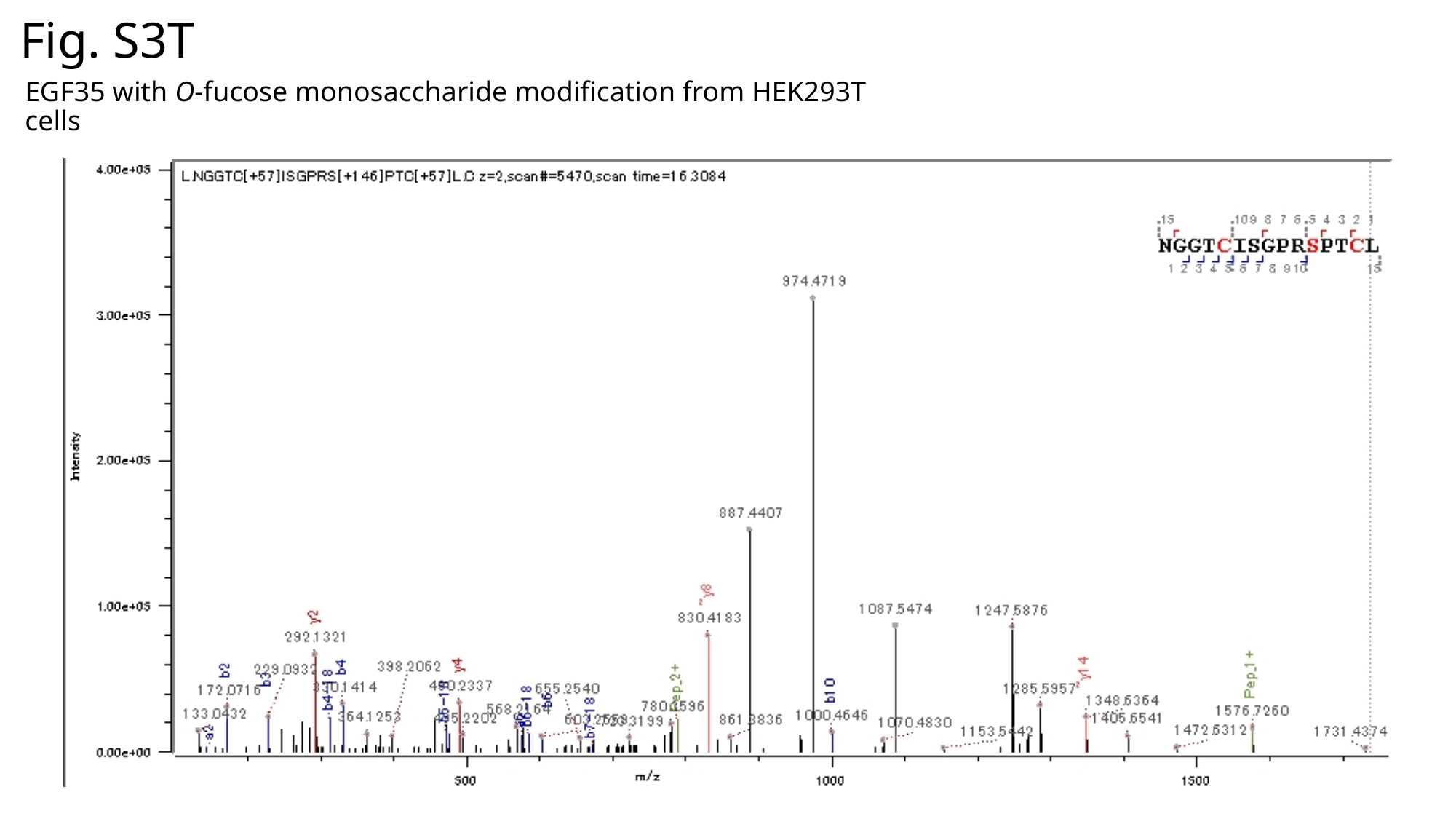

Fig. S3T
EGF35 with O-fucose monosaccharide modification from HEK293T cells

### Slide 23
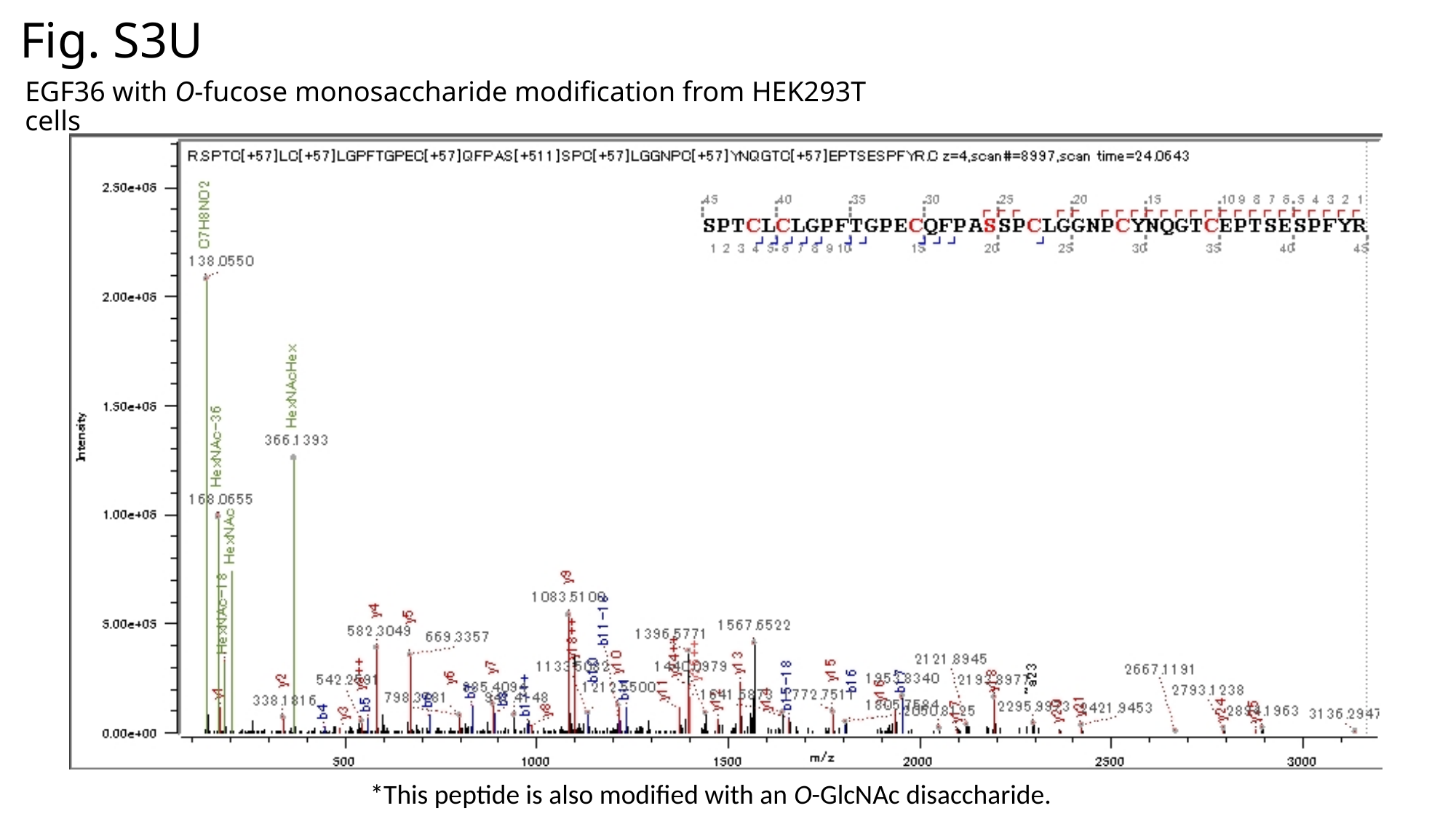

Fig. S3U
EGF36 with O-fucose monosaccharide modification from HEK293T cells
*This peptide is also modified with an O-GlcNAc disaccharide.

### Slide 24
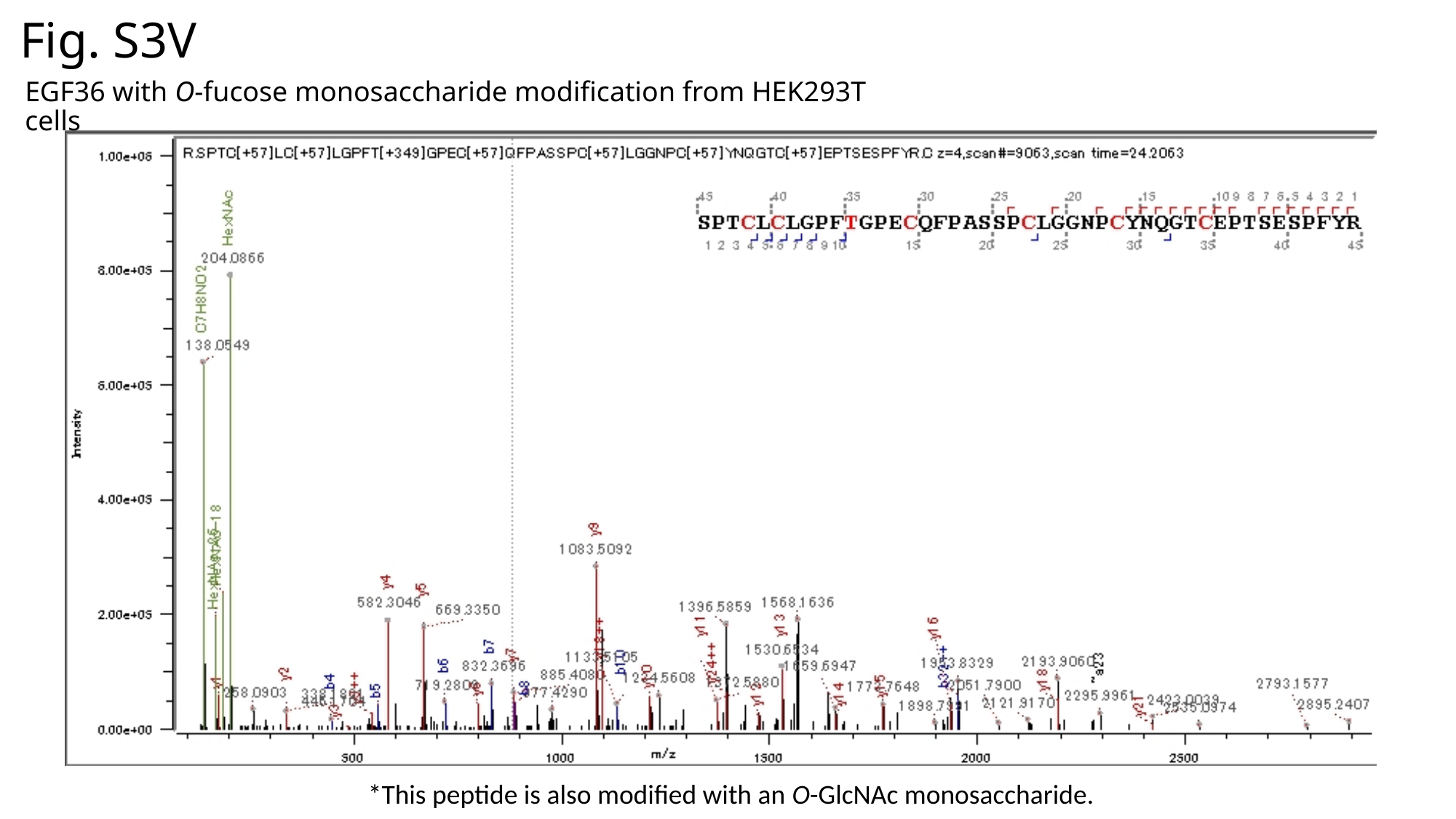

Fig. S3V
EGF36 with O-fucose monosaccharide modification from HEK293T cells
*This peptide is also modified with an O-GlcNAc monosaccharide.

### Slide 25
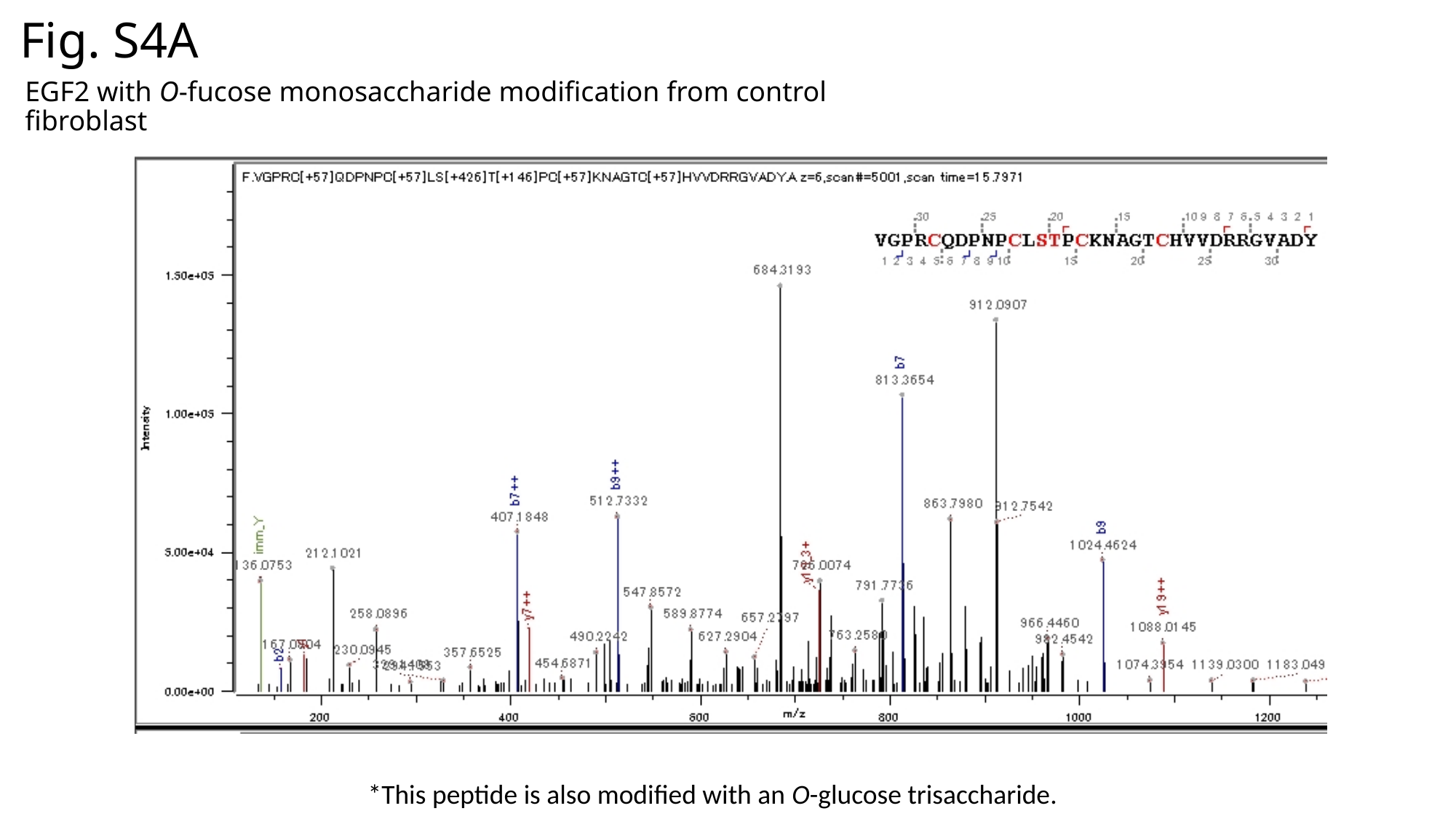

Fig. S4A
EGF2 with O-fucose monosaccharide modification from control fibroblast
*This peptide is also modified with an O-glucose trisaccharide.

### Slide 26
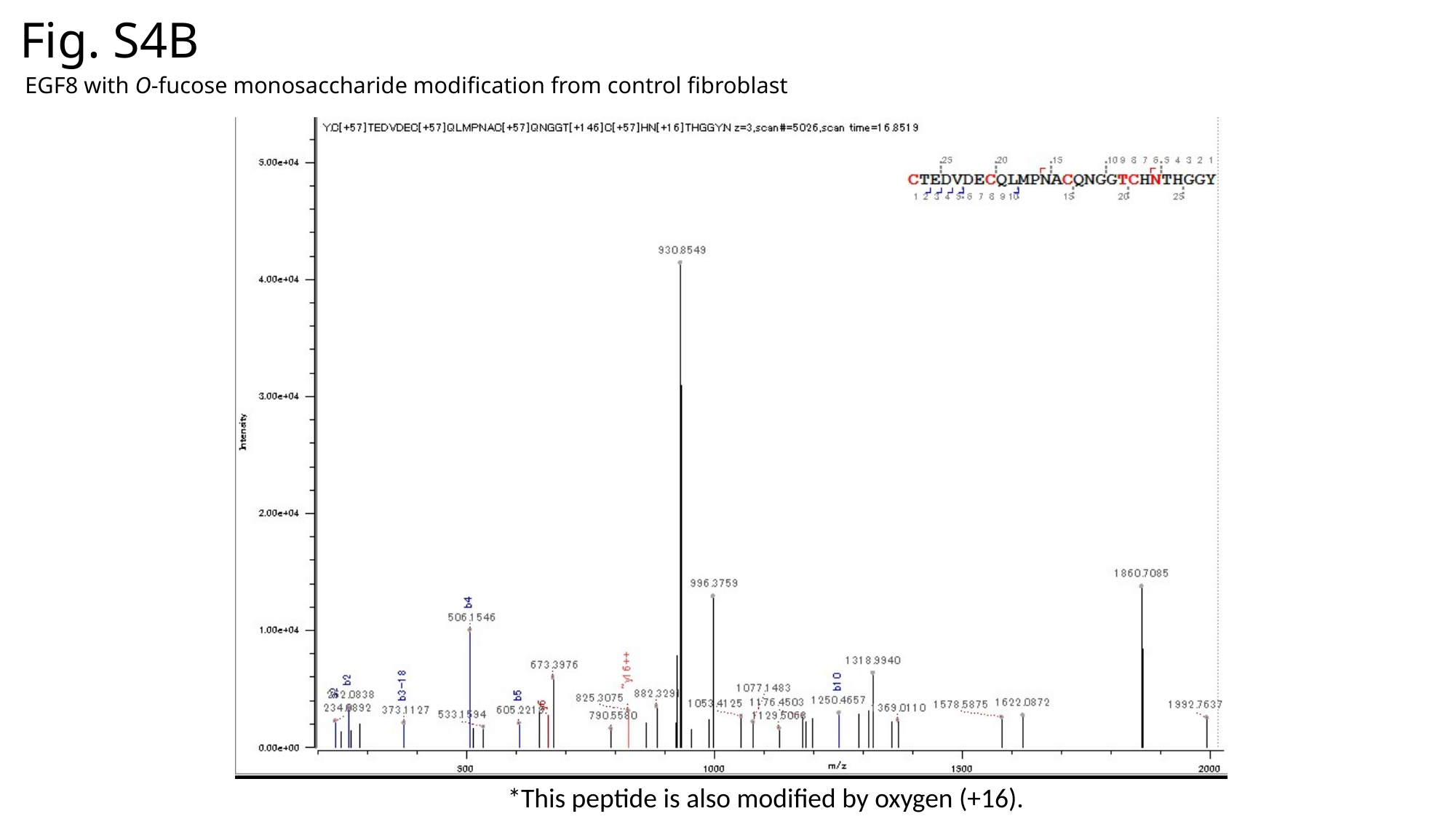

Fig. S4B
EGF8 with O-fucose monosaccharide modification from control fibroblast
*This peptide is also modified by oxygen (+16).

### Slide 27
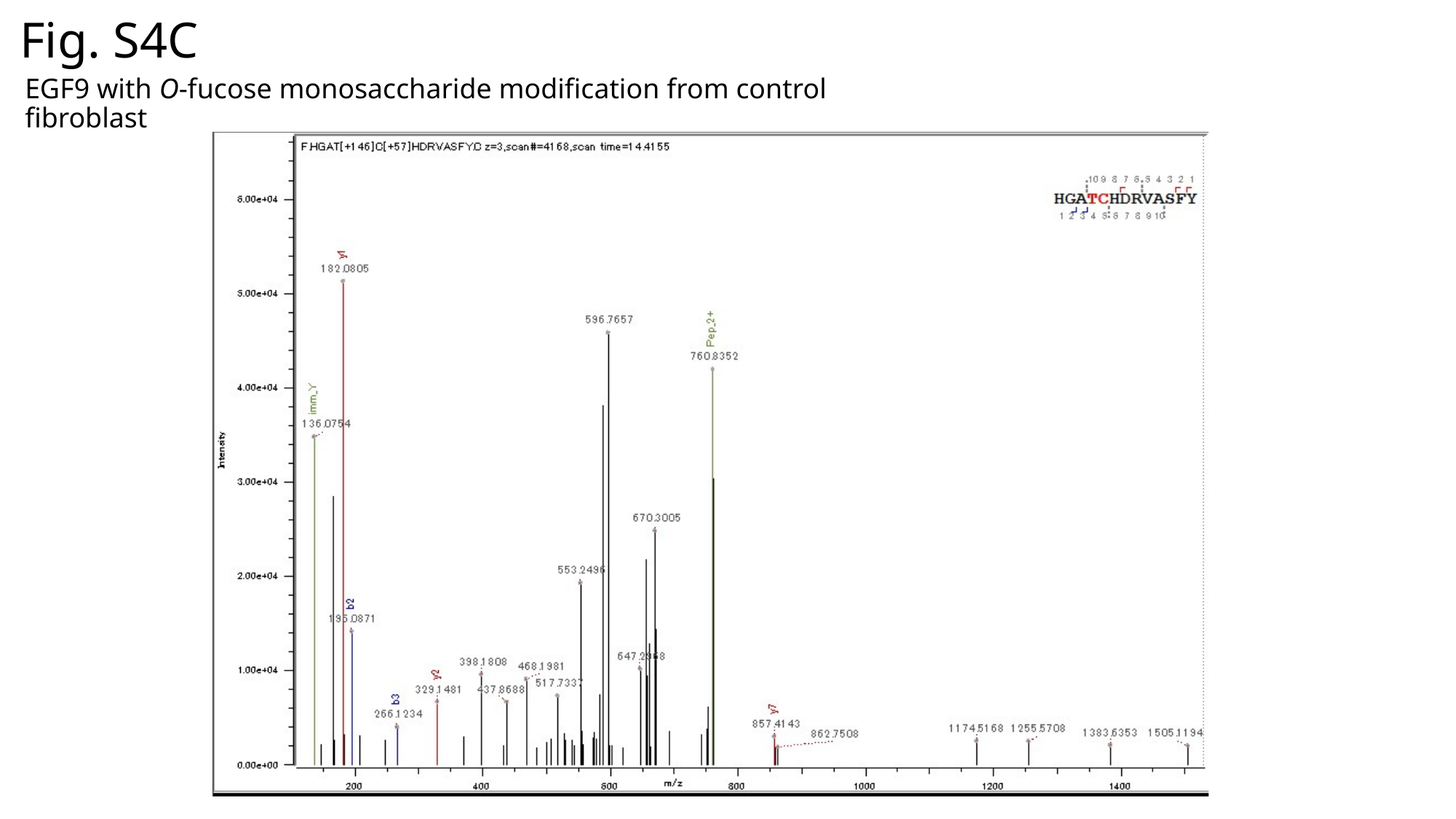

Fig. S4C
EGF9 with O-fucose monosaccharide modification from control fibroblast

### Slide 28
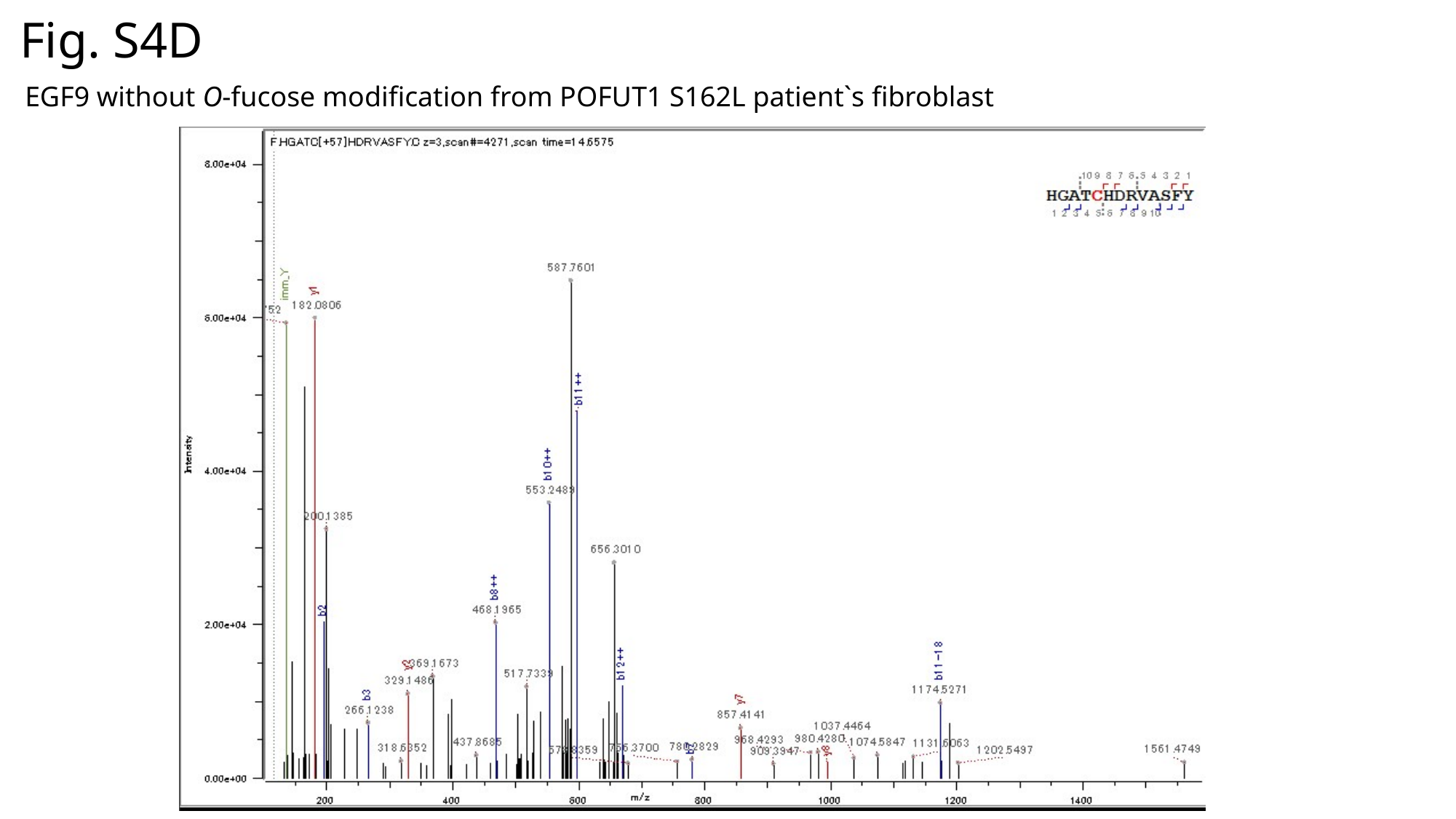

Fig. S4D
EGF9 without O-fucose modification from POFUT1 S162L patient`s fibroblast

### Slide 29
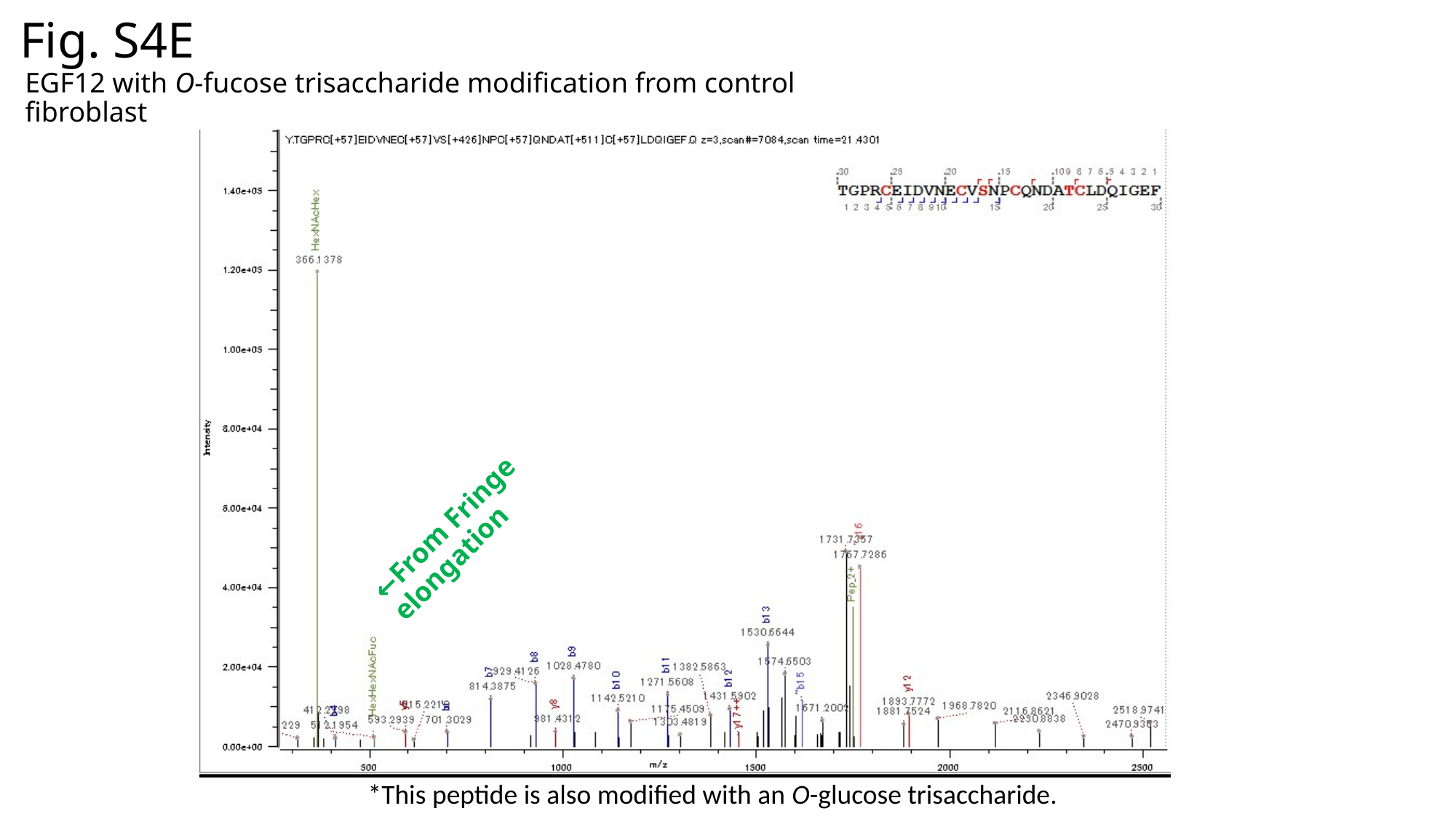

Fig. S4E
EGF12 with O-fucose trisaccharide modification from control fibroblast
←From Fringe elongation
*This peptide is also modified with an O-glucose trisaccharide.

### Slide 30
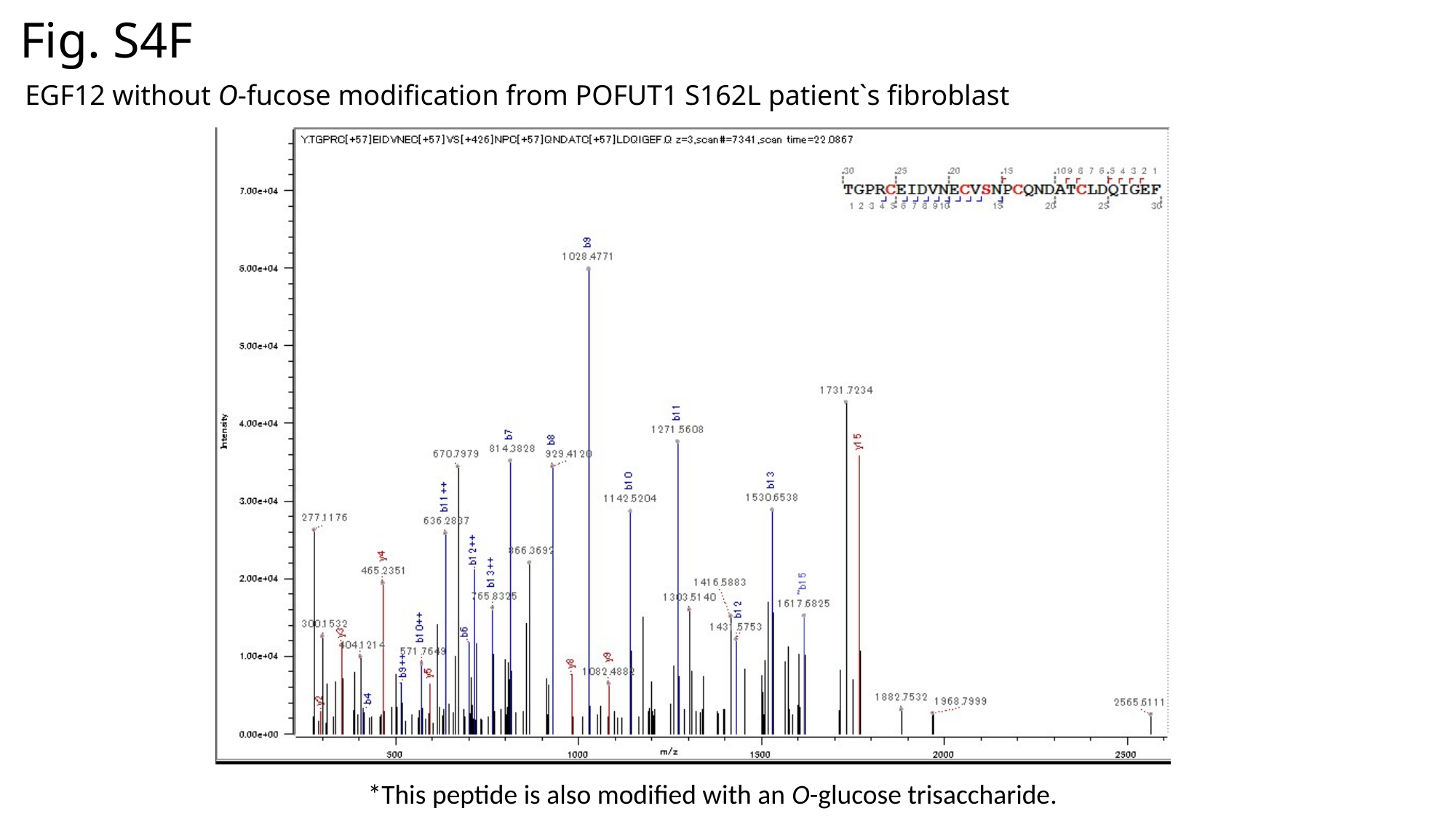

Fig. S4F
EGF12 without O-fucose modification from POFUT1 S162L patient`s fibroblast
*This peptide is also modified with an O-glucose trisaccharide.

### Slide 31
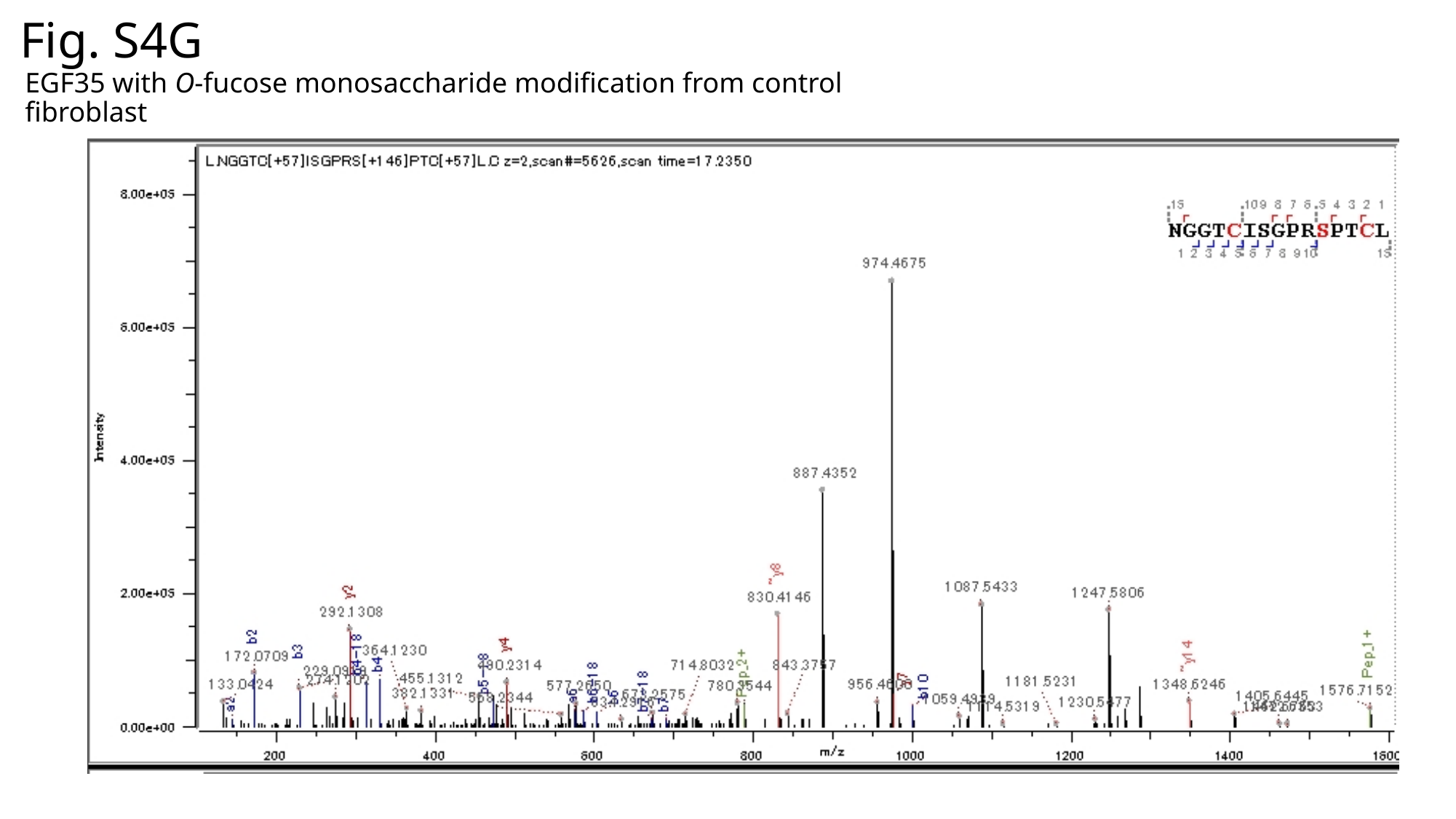

Fig. S4G
EGF35 with O-fucose monosaccharide modification from control fibroblast
